# Hypothermia protects against ventilator-induced lung injury by limiting IL-1β release and NETs formation

**DOI:** 10.1101/2024.09.02.610778

**Authors:** Nobuyuki Nosaka, Vanessa Borges, Daisy Martinon, Timothy R Crother, Moshe Arditi, Kenichi Shimada

## Abstract

Although mechanical ventilation is a critical intervention for acute respiratory distress syndrome (ARDS), it can trigger an IL-1β-associated complication known as ventilator-induced lung injury. In mice, we found that LPS and high-volume ventilation, LPS-HVV, leads to hypoxemia with neutrophil extracellular traps (NETs) formation in the alveoli. Furthermore, *Il1r1^-/-^* LPS-HVV mice did not develop hypoxemia and had reduced NETs, indicating that IL-1R1 signaling is important for NETs formation and hypoxemia. Therapeutic hypothermia (TH) is known to reduce the release of inflammatory mediators. In LPS-HVV mice, TH (32 °C body temperature) prevented hypoxemia development, reducing albumin leakage, IL-1β, gasdermin D (GSDMD) and NETs formation. We also observed that LPS-primed macrophages, when stimulated at 32°C with ATP or nigericin, release less IL-1β associated with reduced GSDMD cleavage. Thus, hypothermia is an important modulating factor in the NLRP3 inflammasome activation, IL-1β release and NETs formation, preventing LPS-HVV-induced acute respiratory failure.

## Introduction

Acute respiratory distress syndrome (ARDS) is a serious pulmonary disorder defined by the onset of non-cardiogenic pulmonary edema and hypoxemia(Force et al. 2012). While it has been more than 50 years since ARDS was first described in the literature, ARDS is still a major cause of respiratory failure in critically ill patients(Matthay et al. 2019). Positive-pressure mechanical ventilation (MV) has become an essential supportive strategy for the management of ARDS(Del Sorbo et al. 2017). However, with the onset of MV as a treatment, it was eventually understood that MV itself can cause and aggravate the lung injury, a condition termed ventilator-induced lung injury (VILI)(Rittayamai and Brochard 2015). Although protective ventilation strategy using low tidal volume has been proven to decrease mortality of ARDS(Pham and Rubenfeld 2017), the mortality rate of ARDS remains as high as 40%(Bellani et al. 2016). Non-injurious MV was reported to be able to still activate proinflammatory signals in the lung(Gharib et al. 2009). Understanding the mechanisms of VILI should lead to novel strategies to further reduce mortality in ARDS(Rittayamai and Brochard 2015).

Neutrophils are key players in the development of ARDS and VILI, as their infiltration is a hallmark of lung injury progression(Grommes and Soehnlein 2011). Over the past decade neutrophil extracellular traps (NETs) have been the subject of intense research with many advances made(Mutua and Gershwin 2021). NETS are a neutrophil-derived meshwork of chromatin fibers decorated with granule peptides and enzymes, and represent a critical host defense strategy against invading microorganisms(Castanheira and Kubes 2019). Intriguingly, recent studies found that NETs play a role in the pathogenesis of tissue injuries, including ARDS and VILI, either with or without infection(Porto and Stein 2016), and NETs have emerged as an important player in COVID-19 pathogenesis(Silva et al. 2022; Veras et al. 2020; Castanheira and Kubes 2023). However, the mechanism of NETs formation and its functional implications in alveolar space during VILI are incompletely understood.

IL-1β has been linked with many inflammatory disorders (Arend and Guthridge 2000). IL-1β levels in BALF were increased in patients with ventilator associated pneumonia (Conway Morris et al. 2010). Furthermore, plasma levels of IL-1β have been associated with worse outcomes in ARDS patients (Meduri et al. 1995). However, while how IL-1β mechanistically affects ARDS is still unknown, it seems that increased amounts of IL-1β is associated with severity of ARDS. NOD-like receptor family pyrin domain-containing protein 3 (NLRP3) inflammasome activation and interleukin (IL)-1β release by alveolar macrophages (AM) was identified as a mechanism for severe acute lung injury (ALI) development in a two-hit model of ARDS using lipopolysaccharide (LPS) instillation and MV(Jones et al. 2014). IL-1β is the most biologically active cytokine in the acute phase of ARDS. It is generally considered that mature IL-1β signals via alveolar epithelial cells resulting in increased lung permeability and pulmonary edema(Ganter et al. 2008). Recent studies have demonstrated that IL-1β and NETs share gasdermin D (GSDMD) as a facilitator of their release(He et al. 2015; Sollberger et al. 2018). Furthermore, IL-1β promotes NETs formation in different experimental settings, which has attracted attention as an uncovered function of IL-1β(Mitroulis et al. 2011; Apostolidou et al. 2016). However, whether IL-1 participates in NETs induction in lung injury is still unknown.

Therapeutic hypothermia (TH) has been generating interest as a promising strategy for ARDS refractory to the current evidence-based therapies(Hayek et al. 2017). TH has long been known to be protective against severe lung injuries clinically and experimentally (Villar and Slutsky 1993; Hayek, et al. 2017). In a porcine two hit model induced by MV and oleic acid, TH reduced the ARDS-associated lung injury and inflammation(Angus et al. 2022). Additionally, one case report showed the successful use of TH for severe refractory hypoxemia in a COVID19 patient(Cruces et al. 2021). This has led to a phase II clinical trial called “cooling to help injured lungs” (CHILL) in which TH is applied in association with neuromuscular blockade in ARDS patients, including those with COVID-19(Shanholtz et al. 2023). TH has strong anti-inflammatory effects and previous studies have reported inhibition of IL-1β production under hypothermia(Diao et al. 2020). However, little is known about how TH affects IL-1β production and NETs formation.

In this study we found that IL-1R1 signaling enhanced NETs formation which contributed to the development of hypoxemia and severe ALI. In addition, we found that TH inhibited IL-1β release from macrophages, which led to less NETs formation and albumin leakage in the alveolar space in our lung injury model. These results add new insights into IL-1β signaling in VILI and ARDS, and its down modulation by TH, which could provide new therapeutic targets.

## Results

### Hypoxemia and NETs formation in alveoli during severe ALI induced by LPS plus mechanical ventilation

To better understand the mechanisms underlying ARDS and VILI, we subjected C57BL/6 mice to intratracheal instillation of LPS and MV with high volume ventilation (HVV) or low volume ventilation (LVV). At 30 and 150 minutes of MV, arterial blood gases were measured, and at the end of 180 minutes, the animals were euthanized and the bronchoalveolar lavage fluid (BALF) was collected (Figure 1A). The combination of LPS and HVV (LPS-HVV) caused prominent reduction in the partial pressure of oxygen (Figure 1B) in the arterial blood (PaO_2_) of mice when compared to the other controls receiving normal saline (NS) and/or LVV instillation, indicative of lung dysfunction. The requirement of both, LPS and HVV for hypoxemia development was confirmed by the strong interaction (p<0.001) between these two factors found by the statistical analysis, and it occurred without significant change in carbon dioxide partial pressure (PaCO_2_), pH, and base excess in the arterial blood (Supplemental Figure 1B-D). Hypoxemia was accompanied by increased neutrophil migration (Figure 1C) compared with NS+HVV without significant change in the number of macrophages (Figure 1D) as determined by flow cytometry (Supplemental Figure 2), and in the total cell number (Supplemental Figure 1E) in BALF. To evaluate the induction of local inflammatory responses in the alveoli, we assessed the levels of several inflammatory markers in BALF. Only LPS, but not HVV, was required for the increase of IL-6, TNFα and CXCL1. On the other hand, only HVV was sufficient to increase the IL-18 release (Supplementary Figure 1, F-I). Although these increased inflammatory mediators do no indicate interactions between LPS and HVV in the proposed model, LPS+HVV increased albumin levels in the BALF (Figure 1E), indicating increased vascular permeability. This injury was associated with increased IL-1β (Figure 1F), IL-1α (Figure 1G), CXCL2 (Figure 1H), plasminogen (Supplemental Figure 1J), and fibrinogen (Supplemental Figure 1K) in the BALF. We next sought to identify if the neutrophils were activated differently between LPS+LVV and LPS-HVV since there was no significant difference in neutrophil migration between the two models (Figure 1C), but only LPS-HVV developed hypoxemia (Figure 1B). We measured the concentration of myeloperoxidase (MPO) (Figure 1I) and neutrophil elastase (NE) (Figure 1J), two neutrophil granule enzymes which are also known to contribute to formation of NETs (McHugh, 2018), as well as the presence of histone-DNA (Figure 1K) and MPO-DNA (Figure 1L) complexes, validated methods for estimating general cell death and NETs(Ishii et al. 2000; Yoo et al. 2014), respectively. We found that LPS-HVV significantly increased all these NETs-associated parameters, indicating that the involvement of NETs in the alveolar space with the local severe acute lung injury resulting with hypoxemia observed in this model.

**Figure 1:**
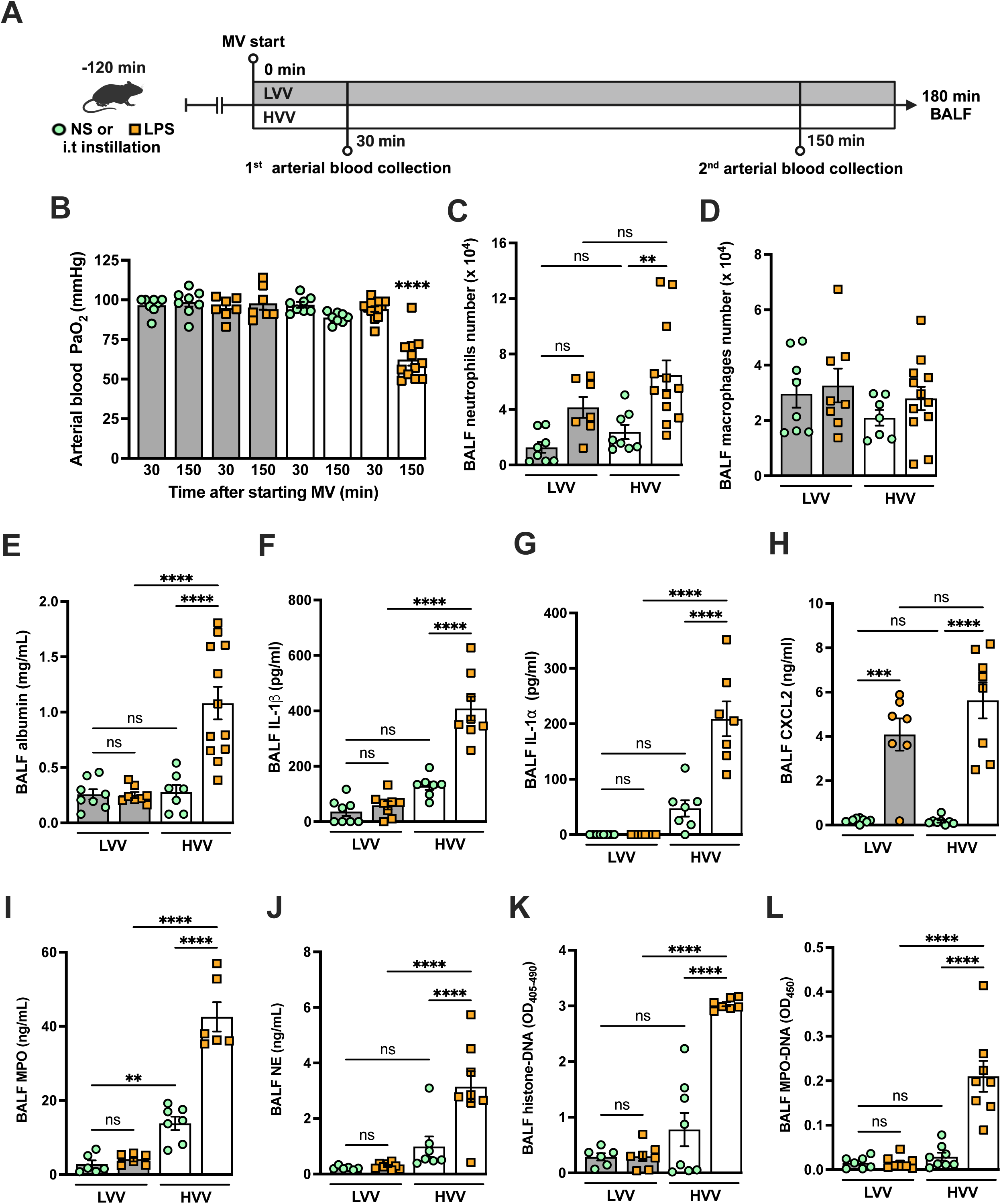
Severe acute lung injury induced by LPS plus high-volume mechanical ventilation is associated with NETs formation in the alveoli. LPS or normal saline (NS) were intratracheally instilled into C57BL/6 mice and, after 120 minutes, the animals were anesthetized and placed on mechanical ventilation (MV) for 180 minutes with the tidal volumes of 30 mL/kg, high-volume ventilation (HVV), or 10 mL/kg low-volume ventilation (LVV) (A). Arterial blood partial pressure of oxygen (PaO_2_) was measured at 30 and 150 minutes after starting MV (B). Absolute counts of neutrophils (C) and macrophages (D) were determined in BALF. The levels of albumin (E), IL-1β (F), IL-1⍺ (G), CXCL2 (H), MPO (I) and NE (J) in BALF collected from euthanized animals after 180 minutes of MV were determined by ELISA. Cell death in BALF was evaluated by measuring histone-DNA complexes (K). NETs formation was evaluated by the detection of MPO-DNA complex (L). ****, ***, and ** indicate *p*<0.0001, *p*<0.001, and *p*<0.01, respectively, determined by three-way ANOVA (B) and two-way ANOVA (C-L) followed by Tukey’s multiple comparisons test; ns, non-significant; the absence of asterisks means non-significant between all the groups; **** on B indicate that the group is different from all the other groups; values are the mean ± SEM; n=7-12.

### Neutrophils are required for the development of severe ALI in the LPS+HVV model

To further investigate the role of neutrophils in this LPS-HVV induced hypoxemia, we depleted neutrophils *in vivo* by treating C57BL/6 mice with antibodies against Ly6G(Carr et al. 2011) eighteen hours before starting MV (Figure 2A). Neutrophil depletion prevented hypoxemia during LPS-HVV, while mice receiving isotype control antibody developed hypoxemia as before (Figure 2B). The absence of neutrophils in the alveoli was confirmed by the reduced total cells number and nearly a complete absence of neutrophils in BALF (Figure 2, C and D). Moreover, in mice depleted of neutrophils, the number of macrophages were quite similar (Figure 2E). The absence of neutrophils significantly reduced the concentration of albumin and IL-6 in BALF (Figure 2, F and H), but did not affect the amount of IL-1β and TNFα (Figure 2, G and I). We also found higher concentrations of the chemokine CXCL2 (Figure 2J) in neutrophil-depleted mice. As expected, neutrophil depletion also resulted in lower MPO and neutrophil elastase (NE) concentrations (Figure 2, K and L), as well as reduced detection of cell death and NETs (Figure 2, M and N) in the BALF.

**Figure 2:**
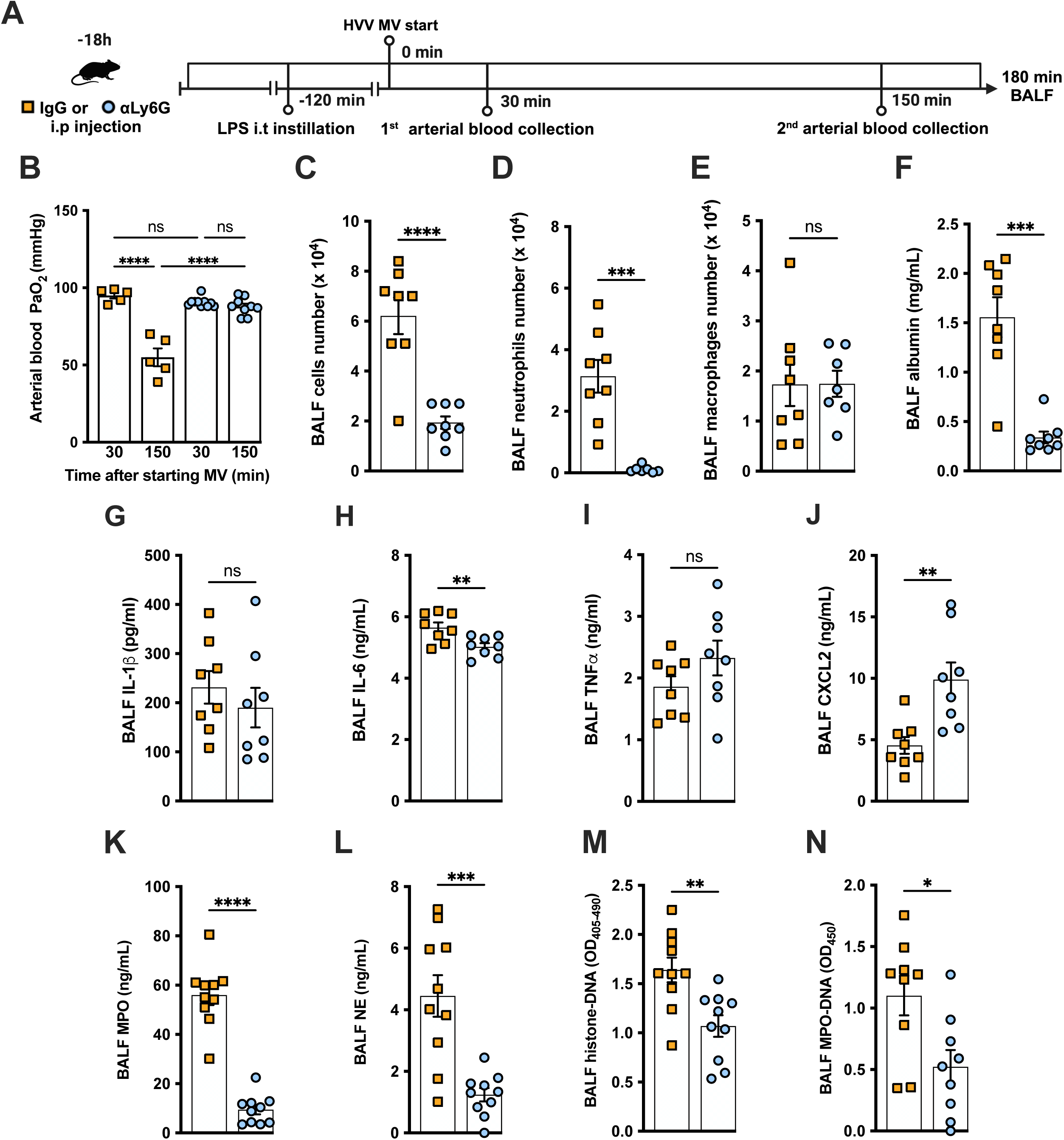
Neutrophils are required for the development of severe acute lung injury in the LPS+HVV model. Eighteen hours before starting mechanical ventilation (MV), the anti-neutrophil monoclonal antibody (αLy6G [1A8]) or the control IgG were administered i.p. to C57BL/6 mice. Sixteen hours later, LPS was instilled i.t. in the mice and, after 120 minutes, they were anesthetized and placed on HVV for 180 minutes (A). Arterial blood partial pressure of oxygen (PaO_2_) was measured at 30 and 150 minutes after starting MV (B). Absolute counts of total cells (C), neutrophils (D) and macrophages (E) in BALF. The concentration of albumin (F), IL-1β (G), IL-6 (H), TNF⍺ (I), CXCL2 (J), MPO (K) and NE (L) were measured in the BALF by ELISA. Cell death and NETs formation in the BALF were evaluated by histone-DNA (M), and MPO-DNA (N) respectively. ****, ***, **, and * indicate *p*<0.0001, *p*<0.001, *p*<0.01, and *p*<0.05, respectively, determined by two-way ANOVA followed by Tukey’s multiple comparisons test (B), unpaired two-tailed Student’s t test (C, E, G-N) or Mann-Whitney test (D, F); ns, non-significant; values are the mean ± SEM; n=5-12.

### NETs contribute to the development of severe ALI in the LPS-HVV model

To investigate the functional role of NETs in the developing hypoxemia in the LPS-HVV-induced severe ALI, we used two different approaches. In the first, we used neutrophil specific PAD4 deficient (*Padi4*^τι/τι^ *S100A8*^cre^) mice and controls (*Padi4*^fl/fl^) (Figure 3A). PADs are required for NETs formation(Rohrbach et al. 2012). Neutrophil specific PAD4 deletion significantly inhibited the hypoxemia development compared with control mice (Figure 3B) but without affecting the numbers of neutrophils (Figure 3C) and macrophages (Supplemental Figure 3B) in the alveoli. It also resulted in reduced levels of BALF albumin but did not change the IL-1β amount in the BALF (Figure 3, D and E). PAD4 deficiency in neutrophils did not alter the levels of MPO and NE in BALF (Supplemental Figure 3, C and D) but did reduce the cell death as measured by histone DNA (Supplemental Figure 3E) and NETs formation as measured by MPO-DNA complexes (Figure 3F). We next treated C57BL/6 mice with DNase I (Figure 3G) aiming to eliminate NETs structures(Czaikoski et al. 2016). This intervention also significantly prevented hypoxemia (Figure 3H) with no impact on neutrophil migration (Figure 3I) and number of macrophage (Supplemental Figure 3G). It also reduced the albumin leakage (Figure 3J) without altering IL-1β levels (Figure 3K) in the alveoli. We observed that DNase I treatment resulted in similar levels of MPO and NE (Supplemental Figure 3, H and I) and resulted in efficiently reduced cell death as measured by histone DNA (Supplemental Figure 3J) and NETs formation as measured by MPO-DNA complexes (Figure 3L).

**Figure 3.**
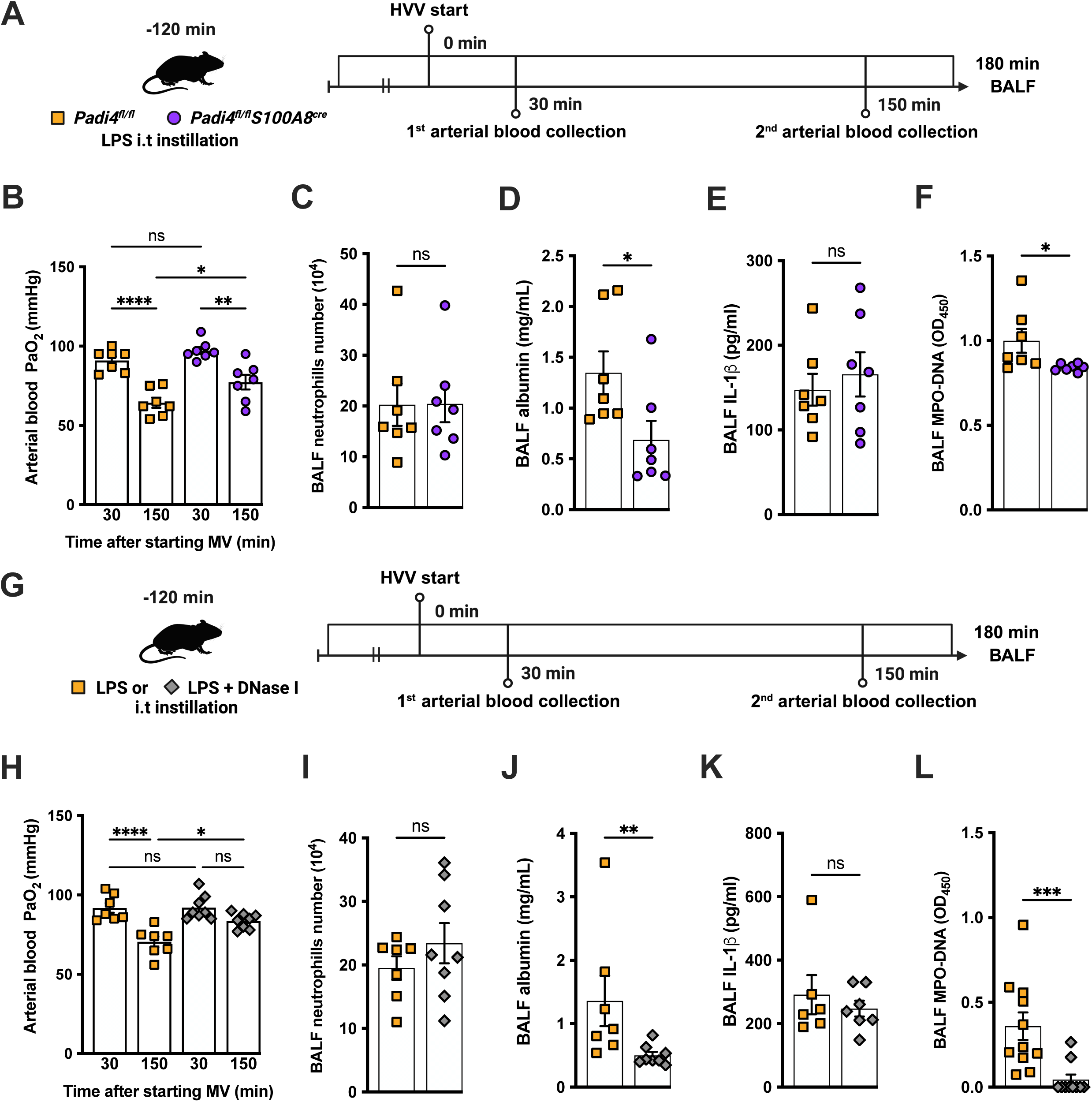
NETs contribute to the development of severe hypoxemia in the LPS-HVV-induced ALI. LPS was instilled to neutrophil specific PAD4 deficient (*Padi4*D^τι/τι^ *S100A8*cre) or the controls (*Padi4*^fl/fl^) mice and, after 120 minutes, the animals were placed on mechanical ventilation (MV) for 180 minutes (A). Arterial blood partial pressure of oxygen (PaO_2_) was measured at 30 and 150 minutes after starting MV (B). Absolute counts of neutrophils (C) and the levels of albumin (D) and IL-1β (E) were measured in BALF. NETs formations in BALF were evaluated by measuring MPO-DNA (F). LPS and DNase I were instilled i.t. to C57BL/6 mice, mice were placed on MV (G). PaO_2_ was measured at 30 and 150 minutes after starting MV (H). Neutrophils (I), albumin (J), IL-1β (K), and MPO-DNA (L) were measured in BALF. ****, ***, **, and * indicate *p*<0.0001, *p*<0.001, *p*<0.01, and *p*<0.05, respectively, determined by two-way ANOVA followed by Tukey’s multiple comparisons test (B, H), unpaired two-tailed Student’s t test (C, E, F, I) or Mann-Whitney test (D, J-L); ns, non-significant; values are the mean ± SEM; n=7-11.

### IL-1R1 signaling is required for NETs formation and severe ALI development in the LPS-HVV model

We previously reported that the activation of NLRP3 inflammasome and IL-1β are required for LPS and MV-induced two-hit model of ALI in mice(Jones, et al. 2014; Nosaka et al. 2020). To confirm the participation of IL-1 receptor type 1 (IL-1R1) downstream signaling in the severe acute lung injury development in this two-hit model, we submitted wild type (WT) and *Il1r1*^-/-^ mice to LPS-HVV (Figure 4A). *Il1r1*^-/-^ mice did not develop hypoxemia, as shown with preserved PaO_2_ values while WT mice have reduced PaO_2_ (Figure 4B). Although IL1R1 deficiency did not affect the number of neutrophils and macrophages in BALF (Figure 4, C and D), it prevented the development of edema resulting from increased vascular permeability, as measured by albumin in BALF (Figure 4E). As expected, IL-1β levels (Figure 4F) were not affected by loss of IL-1R1, demonstrating that there is no feedback loop required for IL-1β production in this model. Loss of this receptor also reduced the release of IL-6 in the BALF (Figure 4G) but did not alter the levels of TNFα and CXCL2 (Figure 4, H and I). Even though the lack of IL-1R1 did not affect neutrophil migration, it’s loss led to a reduction in BALF MPO and NE levels (Figure 4, J and K), as well as cell death (histone DNA) and the NETs formation as measured by MPO-DNA complexes in the BALF (Figure 4, L and M). IL-1β has been found to promote NETs formation in some studies(Mitroulis, et al. 2011; Meher et al. 2018). To confirm the participation of IL-1β in NETs formation *in vitro*, we used neutrophils from a variety of tissues, including bone-marrow neutrophils (BMN), alveolar neutrophils (AN), circulating neutrophils (CN), and peritoneal neutrophils. The purity of these neutrophils was evaluated by flow cytometry (CD45.2^+^CD11b^+^ Ly6G^+^). Isolated BMN, AN, CN and PN were approximately 88, 99, 86 and 89% pure, respectively (Supplemental Figure 4A). Moreover, AN nuclei look more segmentated than the others (Supplemental Figure 4B), and present higher expression of CD11b and Ly6G (Supplemental Figure 4C-E). Neutrophil quality was tested by the baseline of NETs formation (Supplemental Figure 5A) with no stimulus after 4h of incubation. BMN and AN baseline NETs formation was up to 10%, while CN and PN presented about 20% and 60%, respectively, indicating lower stability under cell culture (Suplemental Figure 5, B and C). We then evaluated the NETs formation in response to IL-1β, LPS and ionomycin by neutrophils under cell culture with concentration-response curves, as well as coestimulation experiments (Figure 5A). With BMN and AN we evaluated the concentration-response with single stimuli and costimulation of LPS or IL-1β with ION (Figure 5B-G). The concentration-response curves were aso evaluated with CN and PN (Supplemental Figure 5, C and D). In all samples, the amounts of LPS and IL-1β used were not sufficient to induce NETs formation. Given the better stability of BMN and AN under cell culture, we used these cells for additional experiments. BMN were more resistant to ionomycin, requiring a concentration of 10 μM to induce a 50% NETs formation response, while AN only required 3 μM (Figure 5C, 5F). The sensitivity difference between BMN and AN NETs formation may be related to their maturity state, as AN have highly segmented nuclei and higher CD11b and Ly6G expressions compared with BMN (Supplemental Figure 4, B-E). As LPS and IL-1β did not induce NETs formation by themselves, we next investigated if they could alter the response to ION. For this, neutrophils were stimulated with 10 μg/mL of LPS or 100 ng/mL of IL-1β, and 10 μM (BMN) or 3 μM (AN) of ION and NETs formation was evaluated. We found that IL-1β, but not LPS, enhanced ION-induced NETs formation of both BMN (Figure 5, C and D) and AN (Figure 5, F and G). These data demonstrate that IL-1 signaling is pivotal for hypoxemia development and can modulate NETs formation in LPS+HVV ALI model.

**Figure 4:**
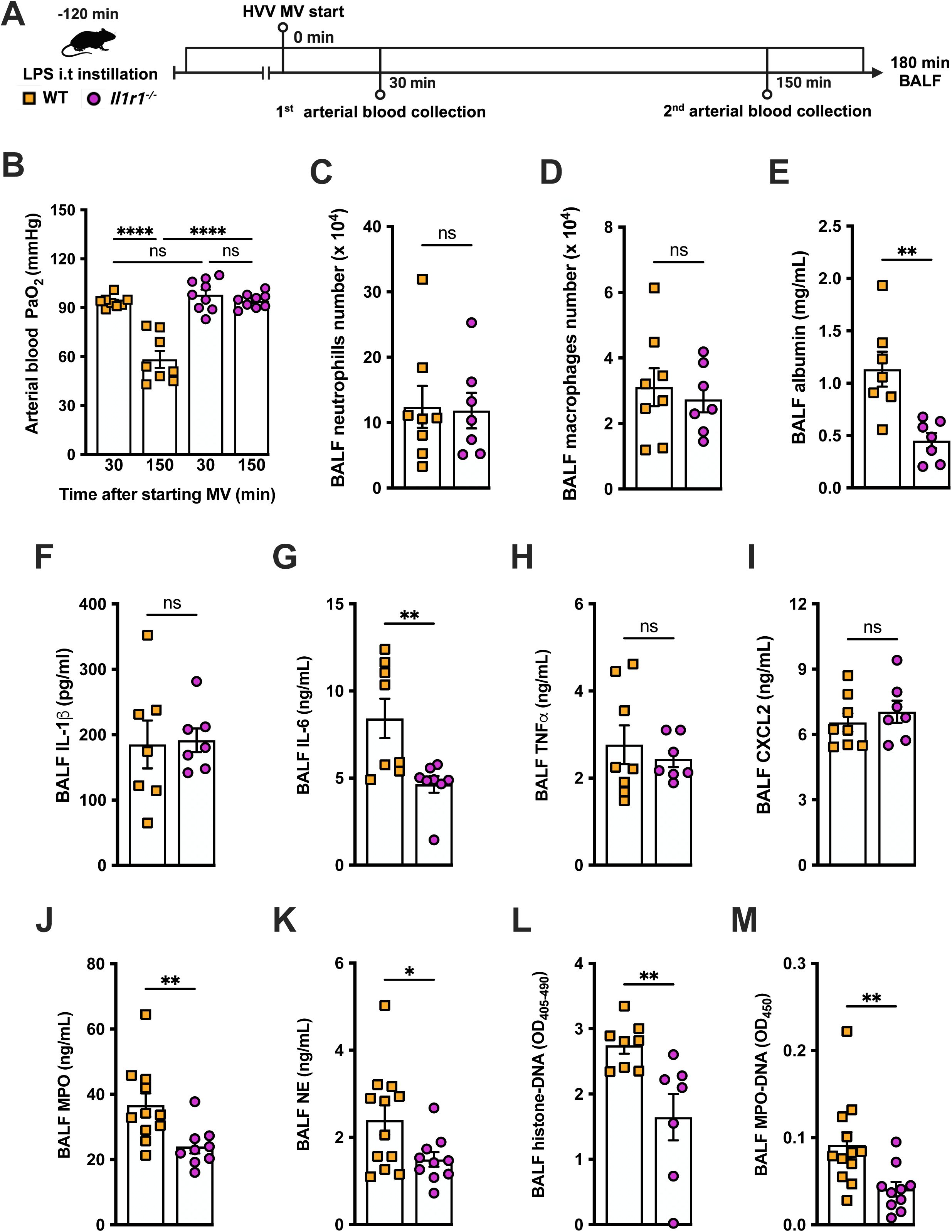
IL-1R1 signaling is required for NETs formation in LPS+HVV-induced ALI. LPS was instilled i.t. into wild type (WT) and *Il1r1^-/-^* mice and, after 120 minutes, the animals were anesthetized and placed on HVV for 180 minutes, followed by sacrifice (A). Arterial blood partial pressure of oxygen was measured at 30 and 150 minutes after starting MV (B). Absolute counts of neutrophils (C) and macrophages (D) in BALF were determined. The levels of albumin (E), IL-1β (F), IL-6 (G), TNF⍺ (H), CXCL2 (I), MPO (J), NE (K) were measured in the bronchoalveolar lavage fluid (BALF) by ELISA. Cell death and NETs formation in the BALF were evaluated by measuring histone-DNA (L), and MPO-DNA (M) respectively. ****, **, and * indicate *p*<0.0001, *p*<0.01, and *p*<0.05, respectively, determined by two-way ANOVA followed by Tukey’s multiple comparisons test (B), unpaired two-tailed Student’s t test (D-F, H-M) or Mann-Whitney test (C, G); ns, non-significant; values are the mean ± SEM; n=7-12.

**Figure 5.**
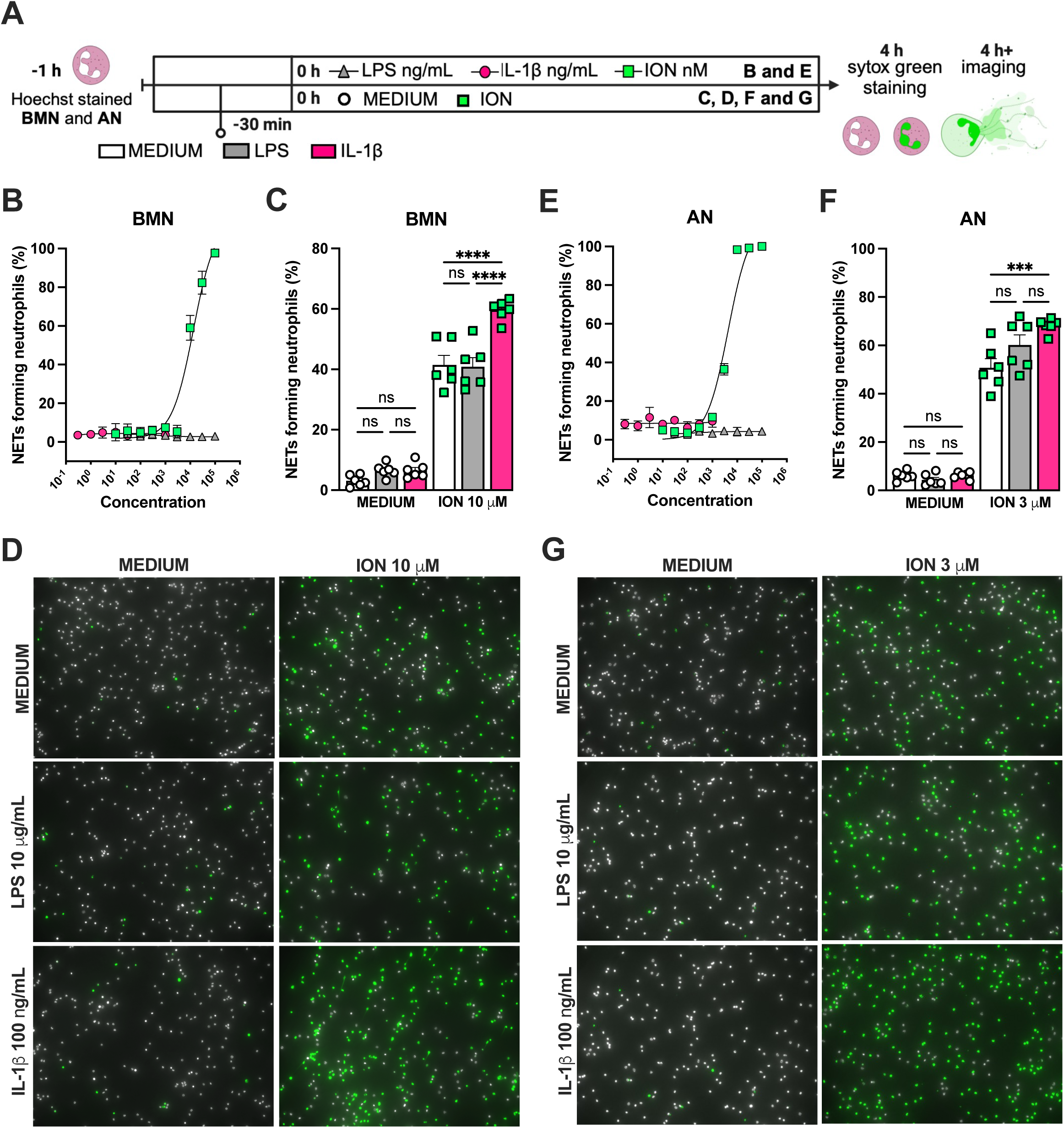
IL-1β enhances ionomycin-induced NETs formation *in vitro.* Hoechst-stained bone marrow neutrophils (BMN) or alveolar neutrophils (AN) were incubated for 1 hour prior to stimulation at 37°C. At time zero the different stimuli were added, and the cells were incubated for 4 hours then stained with Sytox green and the images were captured under microscope (A). BMN (B) and AN (E) were stimulated with several concentrations of lipopolysaccharide (LPS, 30-100,000 ng/mL), IL-1β (0.3-1000 ng/mL), and ionomycin (ION) (10-100,000 nM). For combined stimulation, BMN (C and D) and AN (F and G) were first incubated with LPS or IL-1β 30 min prior ION. The NETs forming neutrophils was analyzed as elongated-shaped Sytox green positive cells and expressed as percentage (%). The Sytox green and Hoechst positive cells are represented by green and white colors, respectively, on the representative images. ****, ***, and ** indicate *p*<0.0001, *p*<0.001 and *p*<0.01, respectively, determined by two-way ANOVA followed by Tukey’s multiple comparisons test; ns, non-significant; values are the mean ± SEM; representative of 3 independent experiments.

### Hypothermia protects against LPS-HVV-induced ALI

During the course of our mouse studies we observed that maintaining normal body temperature was important in obtaining consistent results. Several studies have proven that therapeutic hypothermia can modulate the inflammatory response controlling the release of a variety of inflammatory mediators including IL-1β(Diao, et al. 2020). Thus we decided to investigate whether hypothermia treatement might provide a feasible way to modulate the development of LPS-HVV mediated severe ALI. We then subjected C57BL6 to the LPS+HVV model under controlled body temperature of 37±1 or 32±1 °C, designated as normothermia and hypothermia, respectively (Figure 6, A and B). Hypothermia provided strong protection against hypoxemia, throughout the time course (Figure 6C). While similar neutrophil and macrophage counts were observed between the two groups (Figure 6, D and E), hypothermia resulted in reduced levels of BALF albumin, IL-1β, IL-6, TNFα, MPO and NE (Figure 6, F-K). GSDMD-mediated pore formation impacts not only IL-1β release by macrophages, but was also described to play a role in NETs formation(Sollberger, et al. 2018). We therefore evaluated the soluble GSDMD concentration in the BALF by ELISA (Figure 6L), and found it significantly diminished in mice ventilated under hypothermia. We observed that hypothermia also significantly inhibited the presence of cell death and NETs formation in the BALF in mice subjected to LPS-HVV (Figure 6, M and N). Finally, neigher alkalosis (blood pH > 7.45) nor acidosis (blood pH < 7.35) was observed during hypothermia treatment (Supplemental Figure 6 A and B).

**Figure 6:**
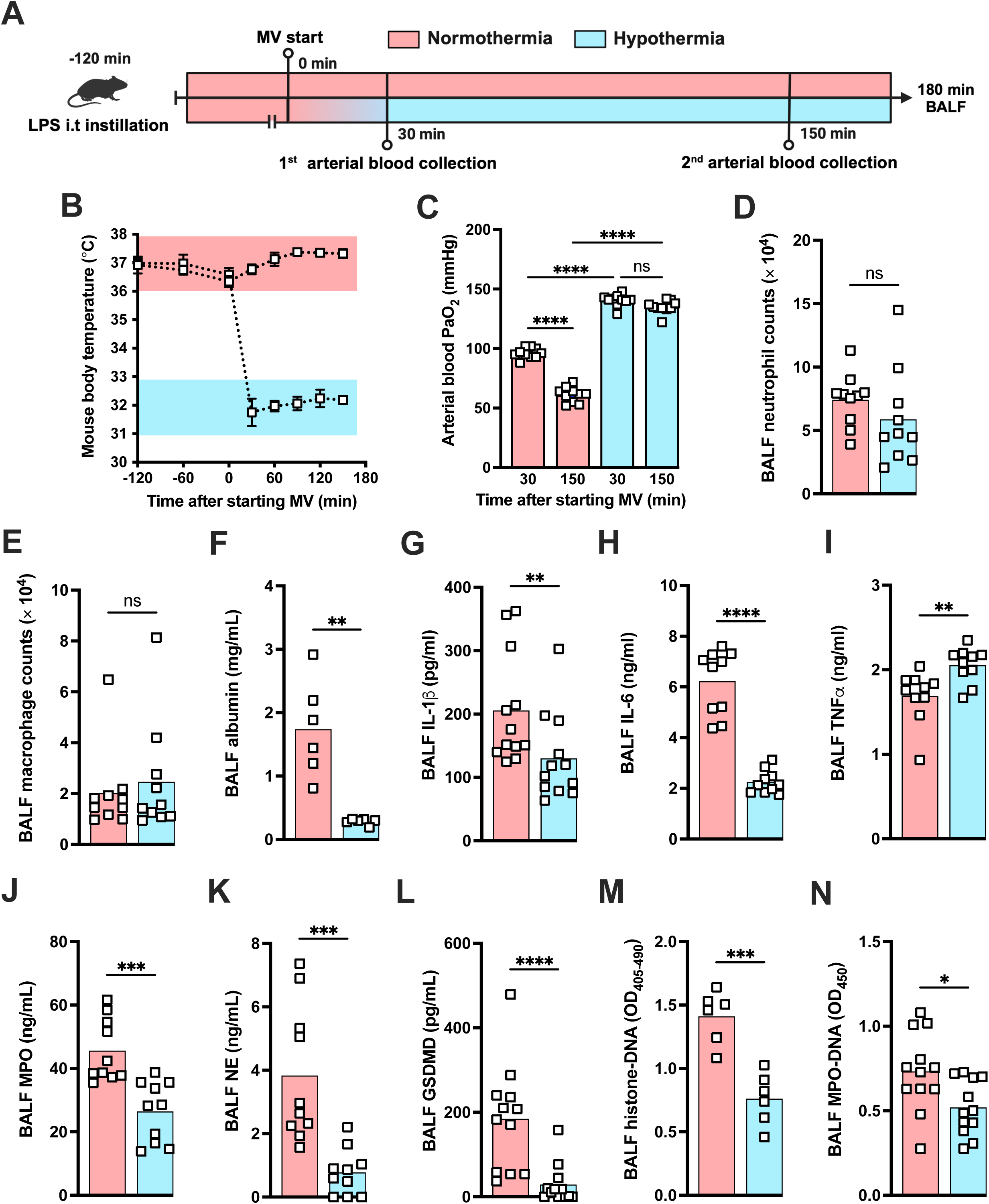
Hypothermia protects against LPS+HVV-induced severe acute lung injury by controlling IL-1β, GSDMD and NETs in the alveoli. LPS was instilled i.t. to C57BL/6 mice and, after 120 minutes, the animals were anesthetized and placed on high-volume ventilation (HVV) for 180 min under controlled body temperature of 37±1 ℃ or 32±1 ℃, designated as normothermia and hypothermia, respectively (A). The body temperature for each group were monitored (B). Arterial blood partial pressure of oxygen was measured at 30 and 150 minutes after starting MV (C). Absolute counts of neutrophils (D) and macrophages (E) in the BALF collected from euthanized after 180 minutes of MV. The levels of albumin (F), IL-1β (G), IL-6 (H), TNF⍺ (I), MPO (J), NE (K) soluble gasdermin D (GSDMD) (L) in the BALF were determined by ELISA. Cell death and NETs formation in the BALF were evaluated by histone-DNA (M), and MPO-DNA (N) respectively. ****, ***, **, and * indicate *p*<0.0001, *p*<0.001, *p*<0.01, and *p*<0.05, respectively, determined by two-way ANOVA followed by Tukey’s multiple comparisons test (C), unpaired two-tailed Student’s t test (D, J, K, M, N) or Mann-Whitney test (E-I, L); ns, non-significant; values are the mean ± SEM, n=6-12 mice/group.

### Hypothermia inhibits macrophage IL-1β release by modulating NLRP3 inflammasome-induced GSDMD cleavage

Since we found that hypothermia inhibited two-hit induced acute respiratory failure with reduced IL-1β in the airways, we next evaluated the ability of bone marrow derived macrophages (BMDMs) to release IL-1β under hypothermia. BMDM were primed with LPS for 3h at 37 ℃, incubated at 37 ℃ or 32 ℃ for 30 minutes prior to adenosine triphosphate (ATP) or nigericin (NIG) treatment and incubated for another 30 min (Figure 7A). Macrophages incubated at 32 ℃ released significantly less IL-1β compared with those incubated at 37 ℃ (Figure 7, B, E). The mechanism by which hypothermia inhibits IL-1β release seems to be independent on caspase-1 activation, as there was no difference in caspase-1 acitvity assay by FLICA between 37 ℃ or 32 ℃ treated macrophages (Figure 7, C and D), but we found that hypothermia resulted in reduced caspase-1 release in the supernatant (Figure 7E). Cleavage of GSDMD is a late limiting step for inflammasome-mediated IL-1β release because its N terminal fragment forms pores on macrophages plasma membrane where the intracellular cytokine crosses into the extracellular compartment(He, et al. 2015). Thus we investigated the effect of 37 ℃ or 32 ℃ temperature on GSDMD expression and cleavage in BMDMs by immunofluorescence, and observed that at 32 ℃ BMDMs express less GSDMD with reduced GSDMD cleavage (Figure 7, F-H). Furthermore, 32 ℃ treatment inhibited GSDMD secretion. These data corroborated our obsevation that while hypothermia inhibited the IL-1β release in the supernatant, there was a concomitant accumulation of unreleased mature IL-1β in the macrophages incubated at 32 ℃ (Figure 7E). Another mechanism by which the NLRP3 inflammasome actitivy is regulated is by autophagy(Nosaka, et al. 2020). To identify whether hypothermia regulates IL-1β release by inducing autophagy, we isolated BMDMs from *Atg16l1^fl/fl^* or *Atg16l^τι/τι^Lysm^Cre+^* mice and repeated the previous experiment. Hypothermic condition (32 ℃ temperature) again significantly inhibited IL-1β release even in macrophages with impaired autophagy, suggesting that the hypothermia effect is independent of autophagy (Supplemental Figure 7), and predominatly through diminished activation of GSDMD.

**Figure 7:**
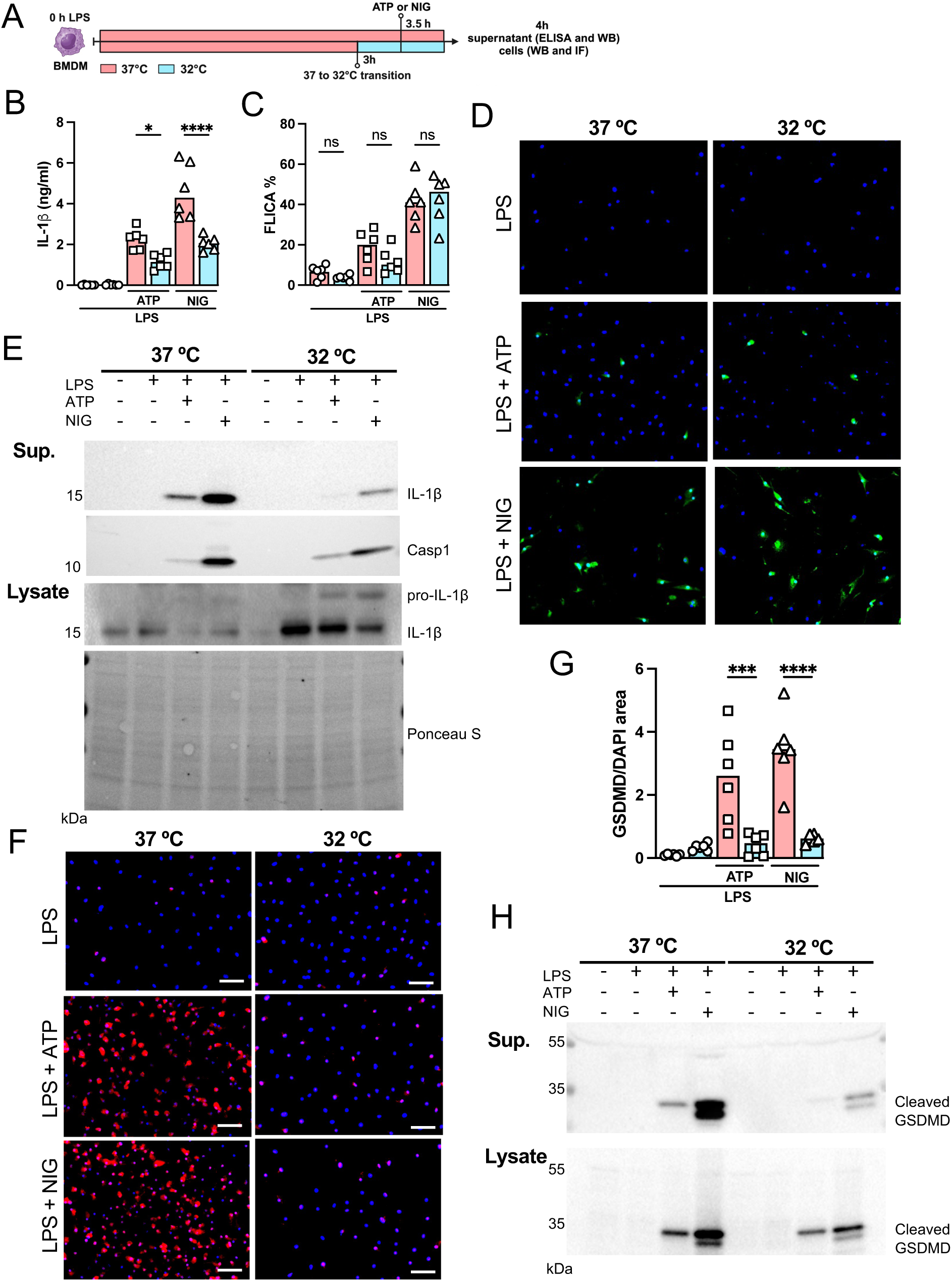
Hypothermia inhibits macrophage IL-1β release by modulating NLRP3 inflammasome-induced gasdermin D cleavage. BMDM were primed with LPS for 3h at 37 ℃, then incubated at 37 ℃ or 32 ℃ for 30 min prior to ATP or nigericin (NIG) treatments for another 30 min (A). In the supernatant, IL-1β concentration was determined by ELISA (B). The supernatant (C) and cell lysate (D) were used for western blotting (WB) analysis, and the resulting membranes were stained for IL-1β, caspase-1 and gasdermin D (GSDMD)(C). WB analysis were also made in the cell lysate, were we investigated the expression of IL-1β and GSDMD. The protein distribution in the cell lysate samples was certified by ponceau staining. BMDM were stained with anti-GSDMD (red) and DAPI (blue), as shown in the representative images. The GSDMD area was analyzed and normalized by DAPI area (E). To evaluate caspase-1 activity, the cells were stained with FAM-YVAD-FMK FLICA and analyzed under the microscope (F). ****, ***, and * indicate *p*<0.0001, *p*<0.001 and *p*<0.05, respectively, determined by two-way ANOVA followed by Tukey’s multiple comparisons test; ns, non-significant; values are the mean ± SEM; representative of 3 independent experiments.

Finally, we investigated the impact of hypothermia on NETs formation *in vitro*. BMN or AN were preconditioned at 37°C or 32°C 1 hour prior to stimulation, and then stimulated for 4 hours with 10 or 3 μM of ION, respectively (Supplemental Figure 8A). We observed significantly reduced NETs formation in both BMN and AN stimulated at 32°C when compared to experiments conducted at 37°C (Supplemental Figure 8, B and C). Taken together, these data demonstrated that hypothermia could be a therapeutic strategy to modulate both IL-1β release and NETs formation for preventing the development of severe acute respiratory failure.

## DISCUSSION

Recent studies suggest that excessive NETs formation plays an important role in the development of multiple diseases, including acute lung injury(Porto and Stein 2016). We have previously reported that NLRP3 inflammasome activation and IL-1β release from alveolar macriohages (AM) are required for the development of hypoxemia in a mouse model of VILI induced by LPS plus mechanichal ventilation, and inhibition of IL-1β signaling via anakinra (IL-1RA), an IL-1 receptor antagonist, attenuates the hypoxemia in this model(Jones, et al. 2014). However, IL-1α also signals through the IL-1R1. Thus is asecond study confirmed that neither anti-IL-1α treated mice nor IL-1α KO mice were protected (Nosaka, et al. 2020). Furthermore, IL-18 is not sufficient to induce Hypoxemia as Saline+HVV treated mice do not develop hypoxemia but still induce IL-18 (Nosaka, et al. 2020). Thus, IL-1β plays a role in inducing hypoxemia during LPS+HVV but neither IL-1α nor IL-18. In the present study, we now show that IL-1β signaling is important for NETs formation in LPS-stimulated mice undergoing mechanical ventilation, with supporting data demonstrating that IL-1β enhances ionomycin-induced NETs formation *in vitro*. Other studies have also reported that IL-1β signaling participates in NETs formation by human neutrophils, that demonstrated that single stimulus with human IL-1β is sufficient to induce NETs(Mitroulis, et al. 2011; Meher, et al. 2018). IL-1β plays a role in the inflammation observed in the lungs of ARDS patients, and the IL-1β level correlates with the severity of disease in these patients(Meduri, et al. 1995; Pugin et al. 1996). Understanding how IL-1β modulates NETs formation in VILI further enhances the importance of IL-1β as a master cytokine in acute lung injury pathophysiology. In a LPS-induced acute lung injury model, NETs and it components can directly injure endothelial and alveolar epithelial cells(Saffarzadeh et al. 2012). In fact, histones, a component of NETs, have been described as damage-associated molecular patterns (DAMPs), and can activate Toll-like receptors (TLR) on a variety of cells, participating in organ injury development(Li et al. 2022). Previous to the discovery of NETs, neutrophils granule components NE and MPO were already found to cause damage to endothelial glycocalyx(Klebanoff, Kinsella, and Wight 1993), which also contribute to lung injury. Neutrophil-derived IL-1β was shown to be released on NETs DNA fibers(Apostolidou, et al. 2016), which could interact directly with the alveolar capillary barrier, leading to an increase in lung epithelial and endothelial permeability(Roux et al. 2005). In addition to the direct role for lung injury, NETs could indirectly aggravate lung injury by inducing further IL-1β production in macrophages(Hu et al. 2017). Furthermore, released MPO on NETs fiber was shown to be active(Parker et al. 2012), which could also activate alveolar macrophages(Grattendick et al. 2002) and lead to additional IL-1β production. Our data support and expand the premise that IL-1β plays a significant role as a driver of the vicious cycle invoved in ventilation-induced acute lung injury and ARDS.

The two-hit acute lung injury models induced by LPS plus MV have been widely accepted and studied extensively(Domscheit et al. 2020). Although MV with 20 ml/kg of tidal volume has been generally accepted as clinically relevant HVV, which we have also used so far, Wilson and colleagues showed that this volume was unlikely to induce substantial lung overstretch in mice(Wilson, Patel, and Takata 2012). In this study, we have slightly modified on our former model, using 30 ml/kg of tidal volume, to have higher volume of MV. Moreover, we added 3 cm H_2_O PEEP to avoid atelectrauma and to focus on injury by lung overstretch, known as volutrauma(Wakabayashi et al. 2014). We confirmed that both LPS and HVV were required for development of acute lung injury in our model, using control mice treated with normal saline and/or LVV. The two-hit requirement in our model resembles the typical clinical ARDS-developing scenario that involves pneumonia and MV, as well as the two-signal for IL-1β production and release. LPS works as the primary signal to induce pro-IL-1β production, and HVV activates NLRP3 inflammasome and induce IL-1β release in mouse AMs(Wu et al. 2013). Indeed, we found that IL-1β was prominently higher in BALF of LPS+HVV mice, and was the main driver of our lung injury model.

Only few studies have been focused on NETs formation as an important player in the pathogenesis of VILI(Rossaint et al. 2014; Yildiz et al. 2015; Li et al. 2017). However, NETs are well described as a mechanism in severe COVID-19 infection(Li et al. 2023) and ARDs pathogenesis(Scozzi et al. 2022). We found that both inhibiting NETs formation by abolishing NETs by DNase I treatment attenuated LPS+HVV induced hypoxemia. Previous reports showed increased NETs in systemic circulation or in the lungs in a single-hit VILI mice model, which was markedly reduced by DNase treatment, resulting in attenuated lung injury(Rossaint, et al. 2014; Li, et al. 2017). In another study, Yildiz and colleagues also investigated NETs formation in the lung tissue of a two-hit VILI mice model(Yildiz, et al. 2015). However, their model differed from ours as the authors used (20 ml/kg versus 30ml/kg) as tidal volume, with no PEEP, for their 4 hours of MV (versus 3 hrs in our study)(Yildiz, et al. 2015). These investigators found slightly increased IL-1β levels, as well as high concentrations of DNA and citrullinated histone-h3, indirect measurement of NETs, in BALF upon LPS and MV, but DNase treatment was not sufficient to inhibit hypoxemia(Yildiz, et al. 2015). They also did not observe an attenuation in NETs formation in reponse to IL-1RA treatment in their model, while we found that LPS+HVV in *IL-1R1^-/-^* mice results in reduced levels of NETs in the alveoli. These observation supports the idea that IL-1β may drive more pathogenic NETs formation in LPS+HVV that leads to severe hypoxemia. Given that the role of neutrophils and types of NETs are still not clear in ALI/ARDS, further studies are clearly warranted.

A striking finding in our study was that hypothermia was protective in a LPS+HVV-mediated mouse model of VILI, preventing severe hypoxemia. Not only did hypothermia treatment in our model inihibit IL-1β release, it also prevented NETs formation. A previous study demonstrated that hypothermia is not protective in a single-hit VILI mouse model, however, it corroborates our data by showing that hypothermia did inhibit IL-1β release in the alveoli without altering neutrophil migration(Faller et al. 2010). However, in a rat model of LPS-induced ALI, hypothermia inhibited neutrophil migration to the alveoli(Lim et al. 2003). Several studies reported lower IL-1β levels in BALF from hypothermia-treated animals subjected to LPS-induced lung injury(Lim, et al. 2003; Hong et al. 2005), which is consistent with our results. Lim and colleagues also reported that hypothermia inhibits LPS-induced nuclear factor κB activation in the lungs and in alveolar macrophages stimulated *ex vivo*(Lim et al. 2004). Another study reported that hypothermia attenuated the expression of caspase-1 in traumatic brain injury in rats, with reduced mature IL-1β and caspase-1 in the cerebral cortex(Tomura et al. 2012). In our study, we found that caspase 1 activity was unaltered in BMDMs cultured under hypothermia, but that IL-1β release was impaired and associated with lower GSDMD expression and cleavage. It has been broadly proposed that the cleaved N-terminus GSDMD can form oligomeric pores in the plasma membrane and play a key role in IL-1β secretion(Zou et al. 2021). While LPS-dependent palmitoylation was proposed as a requirement for N-terminus GSDMD pore formation in macrophages(Balasubramanian et al. 2024) the N-terminus GSDMD plasma membrane translocation and pore formation mechanisms are still poorly understood. Indeed, GSDMD binds to mitochondria outer membrane(Yu et al. 2022; Miao et al. 2023) and nuclei(He et al. 2023) besides plasma membrane. Furthermore, several studies have shown that soluble GSDMD is detected in culture supernatant or body fluids(Karmakar et al. 2020; Nagai et al. 2021; Silva, et al. 2022), but our study may be the first reporting GSDMD detection in the BALF in LPS plus MV induced severe ALI. Supporting this, GSDMD can participate in the host defense by binding to pathogen membranes, potentitally forming cytotoxic pores(Lieberman, Wu, and Kagan 2019). We hypothesized that hypothermia would not affect pro-IL-1β production but inhibit inflammasome activation in macrophages. We found that both immature and mature IL-1β were stuck inside the macrophages with activated caspase-1, associated with a significant reduction in mature GSDMD.

One limitation of our study is that the data includes only young male mice. However, despite thgis limitation, we belive these rsults to be generally applicable as human studies have found that mortality in patients with ARDS does not differ between sexes (Heffernan et al. 2011). Additionally, we induced severe acute lung injury by placing mice on HVV 2 hours afther LPS administraton. At this time point, pulmonary inflammation remains minimal (Dagvadorj et al. 2015). Moreover, LPS induces a febrile response in humans (Ganeshan and Chawla 2017) and can cause either hyperthermia and hypothermia in mice, depending on the dose and ambient temperature (Rudaya et al. 2005). Mice and humans also differ in their sensitivity to LPS (Ganeshan and Chawla 2017), highliting limitations when translating these findings to human sepsis or infection responses. It is also important to distinguish between physiological hypothermia (just below 36°C) and therapeutic hypothermia (typically 32-34°C). While physiological hypothermia is a recognized occurrence in humans with severe infections, it is worth exploring whether therapeutic hypothermia serves as a protective response or if maintaining normothermia to hyperthermia has detrimental effects.

In summary, we developed a two-hit model of severe ALI and ARDS in which both LPS and HVV were required to induce hypoxemia in mice. We demonstrated that IL-1β signaling on neutrophils plays a role in NETs induction, which participates in the development of severe lung injury and hypoxemia. Both IL-1β and NETs have been widely implicated as having a key role in the development of ALI. This research adds specific information about the mechanisms by which severe hypoxemia obeserved in ARDS may be associated with IL-1β effects in lung, and NETs formation. These observations suggest that the inflammasome pathway and its downstream mediators, such as IL-1β and Netosis, may be effective therapeutic targets and could be downmodulated by hypothermia during management of ARDS.

## Materials and methods

### Mice

C57BL/6, *S100A8^Cre^*, *Il1r1*^-/-^, *Lysm*^Cre^ mice on C57BL/6 background were purchased from Jackson Laboratories (Bar Harbor, ME). *Padi4^fl/fl^* mice were provided by Dr. Kelly Mowen (Scripps Research, San Diego, CA, USA). *Atg16l1*^fl/fl^ mice were provided by Dr. Shih (Cedars-Sinai Medical Center, Los Angeles, CA, USA). All *in vivo* experiments were performed in mice at 8-12 weeks of age. All animal studies presented here have been approved by the Institutional Animal Care and Use Committee of the Cedars-Sinai Medical Center. All rodent experimental procedure was conducted under approved Institutional Animal Care and Use Committee protocols.

### LPS plus MV Two-hit acute lung injury model

Male mice were anesthetized with isofluorane (Piramal Healthcare, Bethlehem, PA), and orotracheally intubated with an intravenous catheter (BD Insyte Autoguard, 20GA 1.00 in., Becton Dickinson Infusion Therapy Sytems Inc, Sandy, UT). LPS from *Escherichia coli* (LPS-EB Ultrapure, tlrl-3pelps, Invivogen) was diluted in sterile normal saline (0.9% sodium chloride, NS) to a concentration of 0.1 mg/ml, and 2 microliters per gram of body weight (µl/g) of LPS or NS were administered intrathracheally (i.t.) to mice. Two hours after LPS administration, the mice were intraperitoneally (i.p.) anesthetized with ketamine (Ketaved Ketamine HCl, NDC 50989-161-06, Vedco) and dexmedetomidine (Dextomitor dexmedetomidine hydrochloride, 122692-5, Zoetis) mixture prepared in NS, 50 and 1.0 milligrams per kilogram of body weight (mg/kg), respectively, and orotracheally re-intubated with attention to catheter insertion length, and ventilated using a small animals ventilator system (VentElite, Harvard Apparatus) for 180 minutes with a tidal volume of 10 milliliter per kilogram of body weight (ml/kg) in a respiratory rate (RR) of 150 breaths per minute (bpm), low volume ventilation (LVV), or 30 ml/kg of body weight in a respiratory rate (RR) of 35 breaths per minute, high volume ventilation (HVV), and 3 centimeters of water (cmH_2_O) for positive end-expiratory pressure (PEEP). For hemodynamic support, 500 μl of sterile phosphate buffered saline (PBS) were given to each mouse at the onset of mechanical ventilation (MV). A complementary dose of 25 mg/kg of ketamine and 0.5 mg/kg of dexmedetomidine was administered i.p 90 minutes after starting MV, or earlier as needed. Body temperature was maintained at either 37 ±1 °C or 32 ±1 °C using a heating pad (Heated Hard Pad, Hallowell EMC), and measured rectally by using a temperature probe (08D2, DeltaTrack).

### Arterial blood gas analysis

The arterial blood was collected from anesthetized mice via tail artery by nicking ventral side of the tail with a blade. Approximately 100 µL of whole blood was collected using a Heparinized Micro-Hematocrit Capillary Tube (Fisherbrand). Arterial blood gas was analyzed at 30 and 150 minutes after starting MV using i-STAT1 Analyzer and the i-STAT G3+ Cartridges (Abbott, IL), which provided the values of partial pressure of oxygen (pCO_2_) and partial pressure of carbon monoxide (pCO_2_), in millimeters of mercury (mmHg), pH and base excess as milliequivalent per liter (mEq/L).

### Bronchoalveolar lavage fluid

Bronchoalveolar lavage fluid (BALF) was obtained after 180 minutes of MV with 0.5 mL of cold phosphate-buffered saline (PBS) with 2 mM of EDTA by inserting a standard disposable intravenous catheter (BD Insyte Autoguard, 20GA 1.00 in., Becton Dickinson Infusion Therapy Sytems Inc) into the trachea. A small portion of BALF was stained with ViaStain^TM^ AOPI staining solution, prepared in Cellometer® cell counting chambers and the cells were quantified in the Cellometer® Auto 2000 (Nexcelom Bioscience, Lawrence, MA). The supernatant was isolated for ELISA and the remaining cells were stained with PE anti-mouse Ly6G (50-1276-U100, 1:200, Tonbo Bioscience), FITC anti-human/mouse CD11b (35-0112-U100, 1:200, Tonbo Bioscience), APC anti-mouse F4/80 (20-4801-U100, 1:200, Tonbo Bioscience), violetFluor 450 anti-mouse CD11c (75-0114-U100, 1:200, Tonbo Bioscience) and APC/Cyanine7 anti-mouse CD45.2 (109824, 1:200, Biolegend). The percentage of neutrophils (CD11b^+^ Ly6G^+^) and macrophages (CD11c^+^ F4/80^+^) in the gate of CD45.2^+^ cells were determined by flow cytometry in the *Sony SA3800 spectral cell analyzer* (Sony Biotechnology) and analysed analysed using FlowJo software (FlowJo LLC 10.10.0, Becton Dickinson).

### ELISA measurements

Enzyme-linked immunosorbent assay (ELISA) kits were used for quantifying albumin (Mouse Albumin ELISA kit, 99-134, Bethyl Laboratories), IL-1β (IL-1 beta Mouse Uncoated ELISA Kit, 88-7013-88, Invitrogen), IL-1α (ELISA MAX™ Deluxe Set Mouse IL-1α, 433404, Biolegend), CXCL-2 (Mouse CXCL2/MIP-2 DuoSet ELISA, DY452, R&D Systems), myeloperoxidase (MPO, Mouse Myeloperoxidase DuoSet ELISA, DY3667, R&D Systems), neutrophil elastase (NE, Mouse Neutrophil Elastase/ELA2 DuoSet ELISA, DY4517-05, R&D Systems) cell death (Cell Death Detection ELISA, 11544675001, Roche Life Sciences), IL-6 (Mouse IL-6 ELISA Set, 555240, BD Biosciences), TNF-α (TNF alpha Mouse Uncoated ELISA Kit, 88-7324-88, Invitrogen) CXCL-1 (Mouse CXCL1/KC DuoSet ELISA, DY453, R&D Systems), IL-18 (Mouse IL-18 Matched ELISA Antibody Pair Set, SEK50073, Sino Biological) plasminogen (Mouse PLG/Plasmin/Plasminogen ELISA Kit, LS-F10445, Life Span Biosciences), fibrinogen (Mouse Fibrinogen ELISA Kit, LS-F10440, Life Span Biosciences) and GSDMD (Mouse GSDMD ELISA Kit, ab233627, Abcam). In addition, we developed an ELISA based on MPO associated with DNA as previously describe (Caudrillier et al., 2012) with some modifications. For the capture antibody, 800 ng/mL of anti-MPO capture mAb (Mouse Myeloperoxidase DuoSet ELISA, DY3667, R&D Systems) was coated onto 96-well plates overnight at room temperature (RT). After blocking and washing, 25 μl of BALF was added to the wells with 75 μl incubation buffer (1% BSA/PBS) and incubated for 2 hours at room temperature. After washing, incubation buffer containing a peroxidase-labeled anti-DNA mAb (Cell Death Detection ELISA, 11544675001, Roche Life Sciences, dilution 1:10). The plate was incubated for 1 hour at room temperature. After washing, the peroxidase substrate (TMB) was added and, after 15 minutos at RT in the dark, the reaction was stopped by adding H_2_SO_4_ solution and the absorbance at 450 nm wavelength.

### Neutrophil isolation

Neutrophils were isolated from 8-12 weeks old male WT C57BL/6 mice. The purification was made using two different density gradient protocols made with percoll (Percoll®, GE Healthcare, GE17-0891-01) prepared in Hanks balanced salt solution (HBSS). The percoll gradient 1 (PG1) consists of a three layer gradient made with 75, 57 and 52% of percoll, and the percoll gradient 2 (PG2) consists of two layers, 68 and 52% of percoll. The gradients containing 1 ml of cell suspension on the top were centrifuged for 30 minutes, 1500 x g at RT, with soft acceleration and deceleration. Neutrophils were located above the 75% or 68% percoll layers for PG1 and PG2, respectively. To obtain bone marrow neutrophils (BMN), mice were euthanized, and the rear leg bones were removed and flushed with HBSS with 2 mM of EDTA (HBSS-EDTA). The cells were centrifuged for 5 minutes, 450 x g at RT, and the red blood cells (RBCs) were lysed by adding 10 mL of 0.2% NaCl, gently mixing for 30 seconds. The salt balance was recovered by adding 5 mL of 2.3% NaCl. The cells were centrifuged for 5 minutes, 450 x g at RT, and resuspended in 1 mL of HBSS-EDTA. The cell suspension was carefully added to the PG1, and the BMN isolation was performed as above. For alveolar neutrophils (AN), as described in the two-hit acute lung injury model, mice were anesthetized with isoflurane, orotracheally intubated with intravenous catheter and LPS 0.2 mg/kg was i.t instilled. Three days later, BALF was obtained with 5 mL of HBSS-EDTA divided in 5 lavages with 1 ml of HBSS-EDTA. The cells were centrifuged for 5 minutes, 450 x g at RT, and resuspended at 1 mL of HBSS-EDTA. The cell suspension was carefully added to PG2, and the AN were isolated. Circulating neutrophils (CN) were obtained from the blood collected 6 hours after LPS 0.2 mg/kg i.p injection. The total whole blood was collected by retro-orbital bleeding, in a 5 mL tube containing 2.5 ml of EDTA (2 mg/ml in PBS). The RBCs were lysed by adding the blood (maximum 5 mL, 2.5 ml of blood + 2.5 ml of EDTA) to 30 mL of 0.2% NaCl, gently mixing for 30 seconds. The salt balance was recovered by adding 15 mL of 2.3% NaCl. The lysis step was repeated one time in case remaining RBCs were observed in the pellet formed after 5 minutes of centrigugation, 450 x g at RT. After total elimination of RBCs, the pellet was resuspended in 1 mL of HBSS-EDTA. The cell suspension was added to the PG2, and the CN were purified. For peritoneal neutrophil isolation, mice received i.p injections with 1.5 mL of 3% thioglycolate and, after 16 hours, the animals were euthanized and the peritoneal cavity was washed with 5 mL of HBSS-EDTA. The cells were centrifuged for 5 minutes, 450 x g at RT, and resuspended in 1 mL of HBSS-EDTA. The cell suspension was added to PG2, and the neutrophils were isolated. The cells were stained with ViaStain^TM^ AOPI staining solution, prepared in Cellometer® cell counting chambers, quantified in the Cellometer® Auto 2000 and resuspended in RPMI/neutrophils (RPMI 1640, 10-040-CV, Corning supplemented with 1% of Penicillin-Streptomycin Solution, 30-0002-CL, Corning; 2% of Fetal Bovine Serum, FB-02, Omega Scientific; and 1% of MEM Non-essential Amino Acid Solution, Sigma-Aldrich, M-7145). Neutrophil purity was evaluated by flow cytometry by staining the cells with PE anti-mouse Ly6G (50-1276-U100, 1:200, Tonbo Bioscience), FITC anti-human/mouse CD11b (35-0112-U100, 1:200, Tonbo Bioscience), and APC/Cyanine7 anti-mouse CD45.2 (109824, 1:200, Biolegend). The percentage of neutrophils (CD11b^+^ Ly6G^+^) was determined in the gate of CD45.2^+^ cells, analysed using FlowJo software (FlowJo LLC 10.10.0, Becton Dickinson). The cells were also observed using cytospin (Cytospin 4, Thermo Scientific) prepared slides stained with a rapid staining of bood smear (Hemacolor® Rapid staining of blood smear, 111661, Sigma-Aldrich).

### NETs quantification *in vitro*

2.0 x 10^4^ neutrophils were labeled with 2 µM Hoechst 33342 (Immunochemistry Technology, 639) and seeded in flat bottom 96-well plates. We made three sets of experiments to investigate NETs formation under different conditions, using RPMI/neutrophils to prepare all the treatments. In the first set, after resting the cells for 1 hour at 37 °C, the cells were treated with the following: LPS (30-100,000 ng/mL); IL-1β (Recombinant mouse IL-1β protein, ab259421, Abcam) (0.3-1000 ng/mL); and ionomycin (ION) (Ionomycin calcium salt 10634, Sigma-Aldrich) (10-100,000 nM). In the second set, combined stimulus were given to BMN and AN. The cells were firstly rested at 37 °C for 30 minutes, LPS (10 µg/mL) or IL-1β (100 nM) were added and, after 30 minutes, the cells were treated with the concentration of ION sufficient to induce about 50% of NETs formation, which is 10 µM for BMN and 3 µM for AN. In the third set, BMN and AN were placed at 37 °C or 32 °C for 1 hour, and then treated with ION 10 µM for BMN and 3 µM for AN. In all of them, the cells were incubated for 4 hours at 37 °C or 32 °C, maintaining the initial resting temperature setting in each experiment, to allow NETs induction. The cells were then stained with 5 µM of sytox green (SYTOX™ Green Nucleic Acid Stain, 57020, Invitrogen) prepared in sterile PBS, and centrifuged 5 minutes, 500g RT. The images were obtained on a Keyence BZ-9000 (Keyence Corporation of America) microscope at 20x magnification. The NETs forming neutrophils percentage was evaluated manually, according to the total cell number in the field, obtained by the sum of hoechst (white) and sytox green (green) positive cells counted using the Keyence BioAnalyzer software (Keyence Corporation of America).

### Bone marrow-derived macrophage culture

Bone marrow were obtained from 8-12 weeks old male WT C57BL/6, *Atg16l1^fl/fl^* or *Atg16l^τι/τι^ Lysm^Cre^*mice, and BMDM were differentiated in RPMI/macrophages (RPMI 1640, 10-041-CV, corning; supplemented with 1% of Penicillin-Streptomycin Solution, 30-0002-CL, Corning; 50 μM of 2-mercaptoethanol; and 10% of Fetal Bovine Serum, FB-02, Omega Scientific) containing 15% of L929 cell conditioned medium (LCM) for seven days, supplementing the medium with extra 15% of LCM every 3 days.

Cells were washed with PBS, and non-adherent cells were removed; adherent cells were then collected and seeded in a 96-well plate 1 day before stimulation. BMDM were primed with LPS 1 μg/mL for 3 hr, followed by ATP 5 mM (Adenosine 5′-triphosphate disodium salt, A2383-1G, Sigma-Aldrich) or nigericin 10 μM (Nigericin sodium salt, BML-CA421-0005, Enzo Life Sciences) stimulation for 30 min at either 37 °C or 32 °C, and the supernatants were collected for ELISA measurements and western blotting, and the cells were lysed for westen blotting. For evaluating caspase-1 activity and GSDMD expression by immunofluorescence, LPS-primed BMDM were stimulated with 5 mM ATP for 15 min or 10 μM nigericin for 30 min. For Caspase-1 activity a commercial kit (FAM-FLICA™ Caspase-1 Assay Kit, 98, Immunochemistry Technology) was used. Immunoblots were peformed using antibodies against IL-1β (anti-IL-1 beta antibody, 2 μg/mL, ab9722; Abcam), caspase-1 (recombinant anti-pro caspase-1 + p10 + p12 antibody, 1:1000, ab179515; Abcam) and GSDMD (recombinant anti-GSDMD antibody, 1:1000, ab209845, Abcam). The GSDMD antibody used in the immunoblot was also used for immunofluorescence staining, which was mounted with fluorescence mounting medium with DAPI (Mounting Medium With DAPI, ab104139, Abcam). The FAM-FLICA caspase-1 images and the other immunofluorescence images were obtained on a Keyence BZ-9000 *(Keyence Corporation of America)* microscope at 20x magnification.

## Quantification and statistical analysis

All data were analyzed using Prism 9 (GraphPad Software Inc., La Jolla, CA). Normality within each group was assessed using the Shapiro-Wilk and Kolmogorov-Smirnov tests. For comparisons between two groups, the Mann-Whitney U test was used for non-normally distributed data, while the unpaired Student’s t-test was applied to data that met the assumption of normality. One-way ANOVA (with a single independent factor), two-way ANOVA (with two independent factors), and three-way ANOVA (with three independent factors) were performed for comparisons involving more than two groups, followed by Tukey’s post hoc test. A P-value of less than 0.05 was considered statistically significant.

## Acknowledgments

This work was supported by National Institute of Health (NIH) grant R01-HL130353-01 to KS. The illustrations presented in this work were created by using BioRender, https://BioRender.com/m51x648, https://BioRender.com/y06f775, https://BioRender.com/m67j875.

## Author Contributions

K.S. conceived and led the project. N.N, V.F.B, D.M and K.S performed the experiments. N.N, V.F.B and K.S. wrote the manuscript. T.C and M.A provided critical editing and content to the manuscript and experimental design. All the authors read and approved the final manuscript.

## Supplemental information

Suplemmental material. Supplemental Figure 1-8.

