## Supplemental Figure for "Hypothermia protects against ventilator-induced lung injury by limiting IL-1β release and NETs formation"

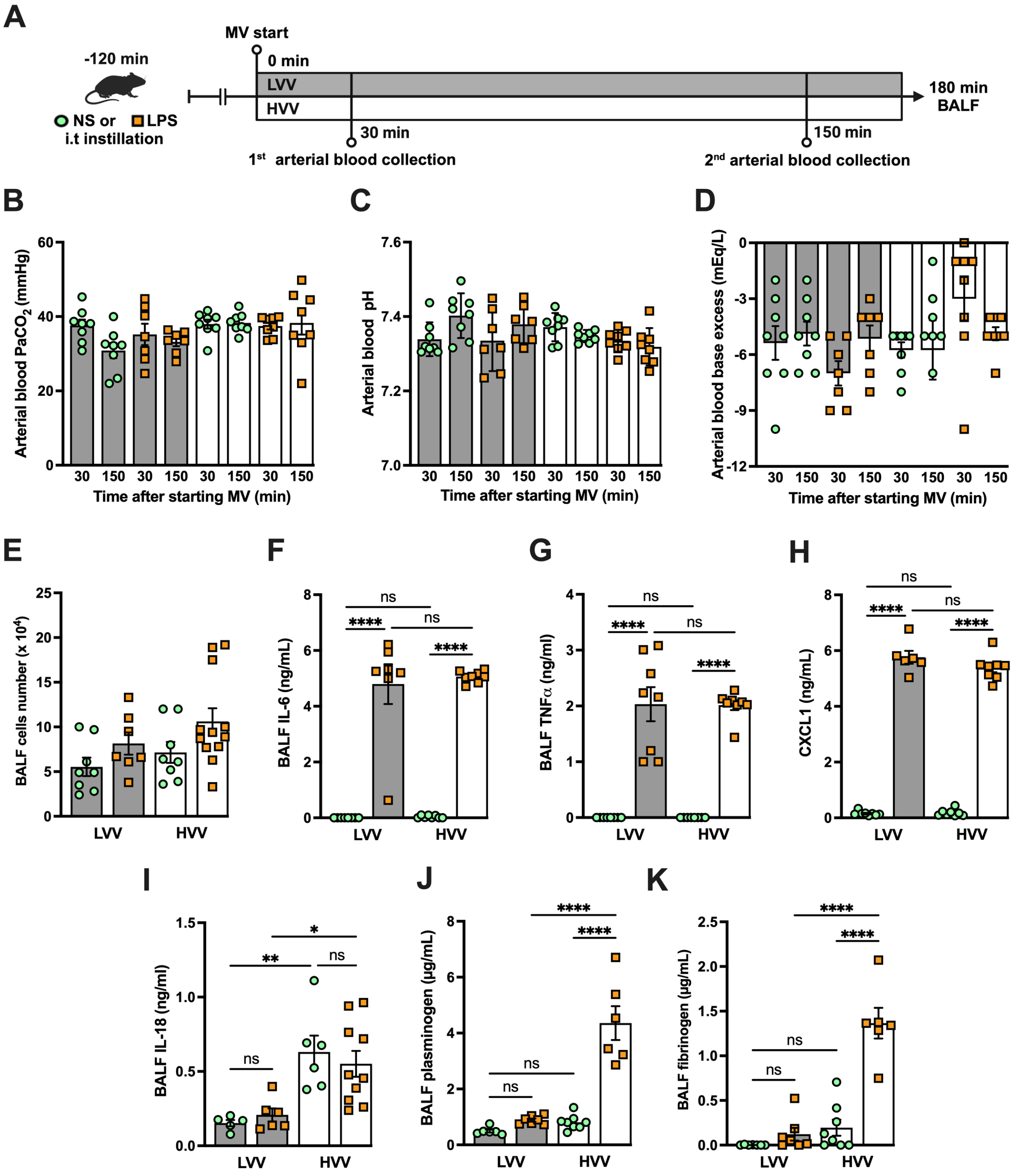


**Supplemental Figure 1 (related to Figure 1): Severe acute lung injury induced by LPS plus high-volume mechanical ventilation occurs without alteration in respiratory acidosis or alkalosis and increased plasminogen and fibrinogen levels in the alveoli**. LPS or normal saline (NS) were i.t. instilled to C57BL/6 mice and, after 120 minutes, the animals were anesthetized and placed on mechanical ventilation (MV) for 180 minutes with the tidal volumes of 30 mL/kg, high-volume ventilation (HVV), or 10 mL/kg low-volume ventilation (LVV) (A). Arterial blood partial pressure of carbon dioxide (arterial blood $PaCO_2$) (B), pH (C) and base excess (D) were measured at 30 and 150 minutes after starting MV. Absolute cell counts (E), IL-6 (F), TNF-α (G), CXCL1 (H), IL-18 (I), plasminogen (I) and fibrinogen (K) were quantified in the BALF collected from euthanized animals after 180 minutes of MV. ****, **, and * indicate $p<0.0001$, $p<0.01$, and $p<0.05$, respectively, determined by three-way ANOVA (B-D) and two-way ANOVA (E-K) followed by Tukey's multiple comparisons test; ns, non-significant; the absence of asterisks means non-significant between all the groups; values are the mean ± SEM; n=7-12.

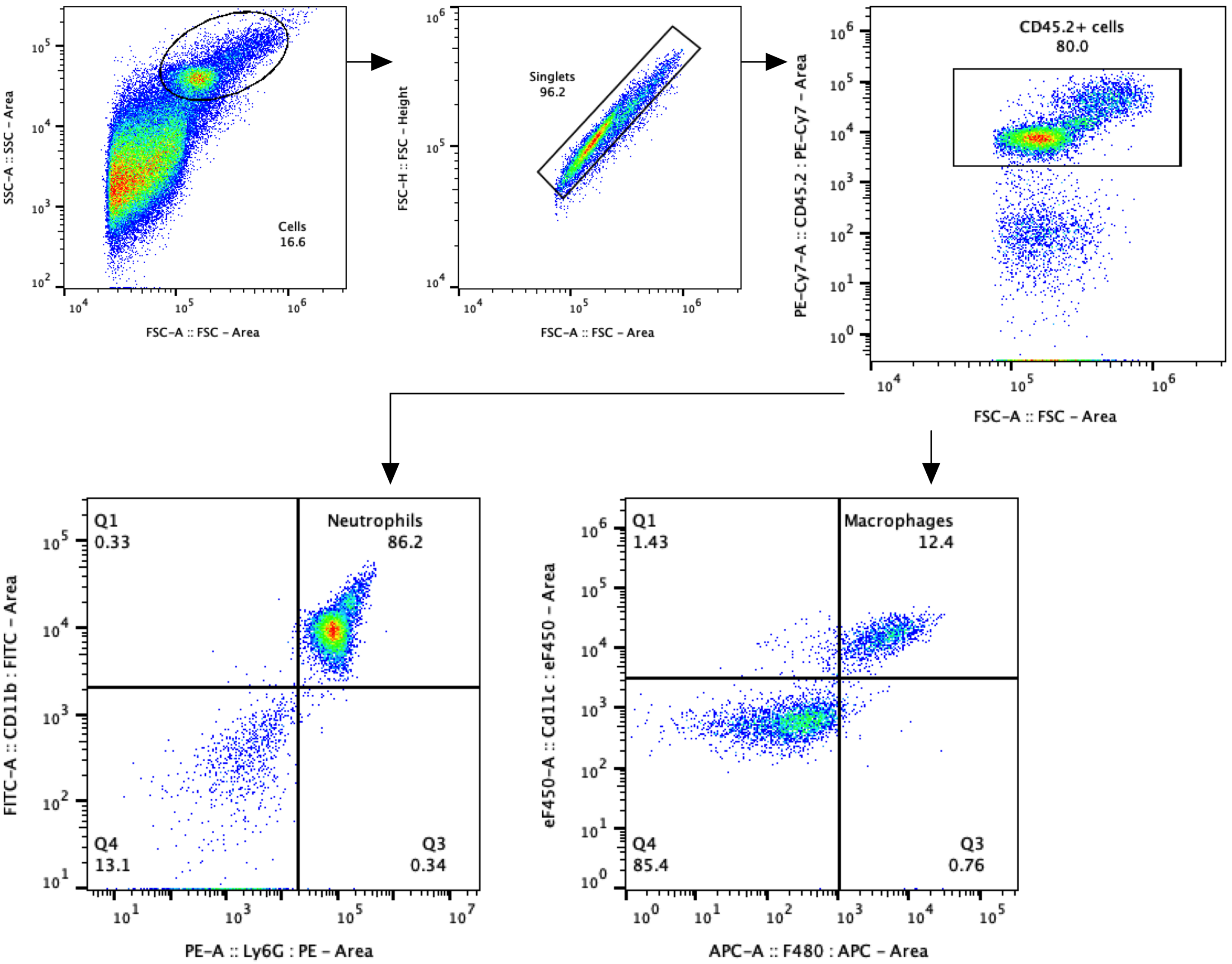


**Supplemental Figure 2 (related to Figure 1): Gate strategy for alveolar neutrophils and macrophages in the two-hit model.**

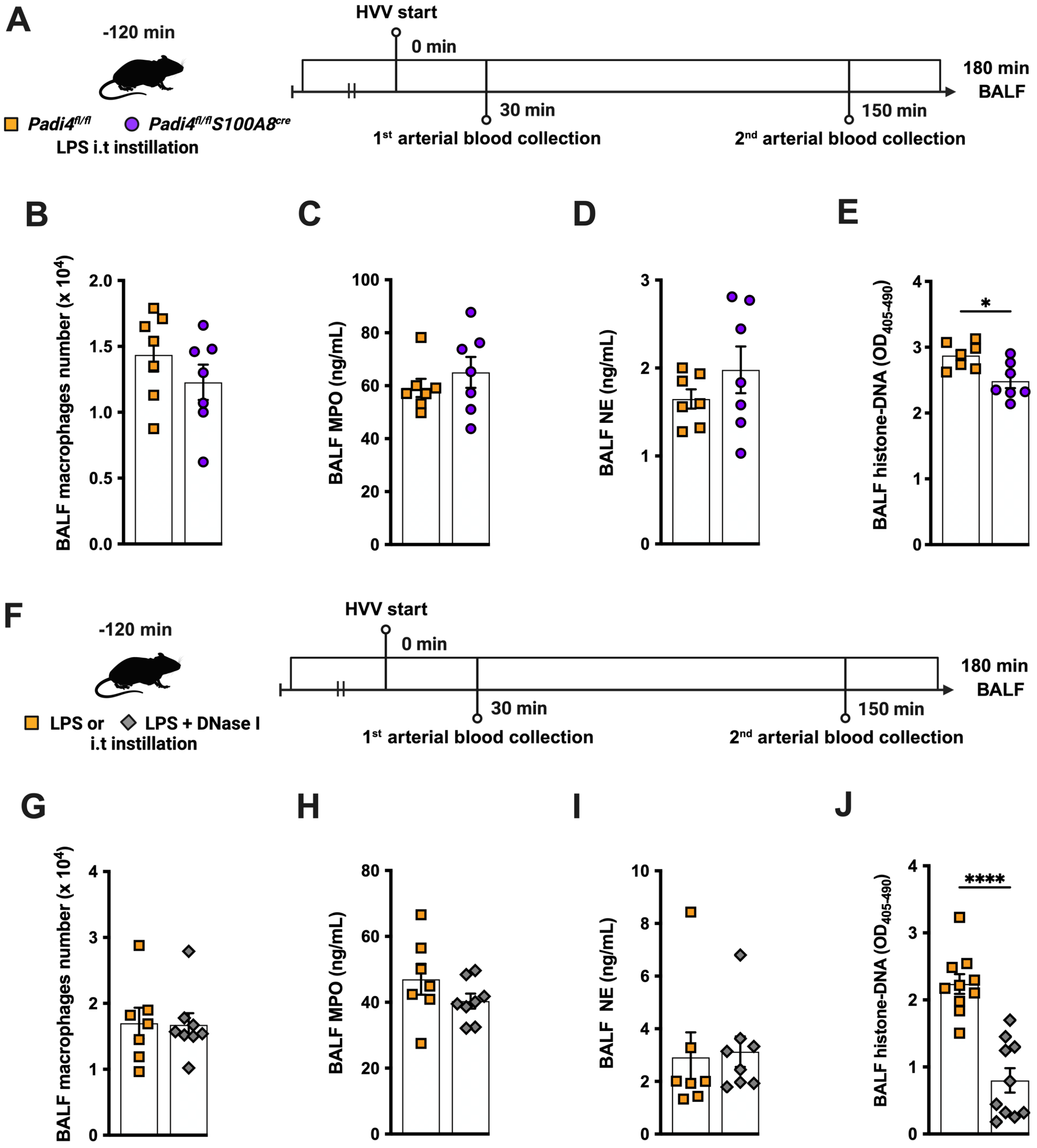


**Supplemental Figure 3 (related to Figure 3): In the LPS-HVV model, neutrophil-specific *Padi4* deletion and DNase 1 treatment reduce the levels of histone-DNA complexes in the alveoli without altering the macrophages number and the levels of myeloperoxidase and neutrophil elastase.** Two different approaches to mitigate NETs effects in the in the LPS-HVV model are presented. In the first (A), we used neutrophil specific PAD4 deficient mice (*Padi4*$^{\Delta/\Delta}$ *S100A8*$^{cre}$) or the controls (*Padi4*$^{fl/fl}$) and the number of macrophages (B), the levels of myeloperoxidase (C) and neutrophil elastase (D), as well as histone-DNA complexes were assessed in the bronchoalveolar lavage fluid (BALF). In the second, simultaneously with the lps instilation, the animals also received DNase I (F) and the number of macrophages (B), the levels of myeloperoxidase (C) and neutrophil elastase (D), as well as histone-DNA complexes were assessed in the BALF. **** and * indicate $p$<0.0001 and $p$<0.05, respectively, determined by unpaired two-tailed Student's t test (B, D-H, J) or Mann-Whitney test (C, I); values are the mean ± SEM; n=7-8.

**A**

| | BMN (n=3) | AN (n=3) | CN (n=3) | PN (n=3) |
|---|---|---|---|---|
| **Cell number (x $10^6$)** | 16.93 ± 1.617 | 4.047 ± 0.169 | 2.95 ± 0.8003 | 2.1 ± 0.2082 |
| **Viability (%)** | 95.3 ± 1.504 | 92.03 ± 0.2186 | 90.57 ± 1.49 | 73.4 ± 2.831 |
| **$CD11b^+Ly6G^+$ (%)** | 88.2 ± 2.787 | 98.8 ± 0.533 | 86.03 ± 2.184 | 89.4 ± 1.55 |

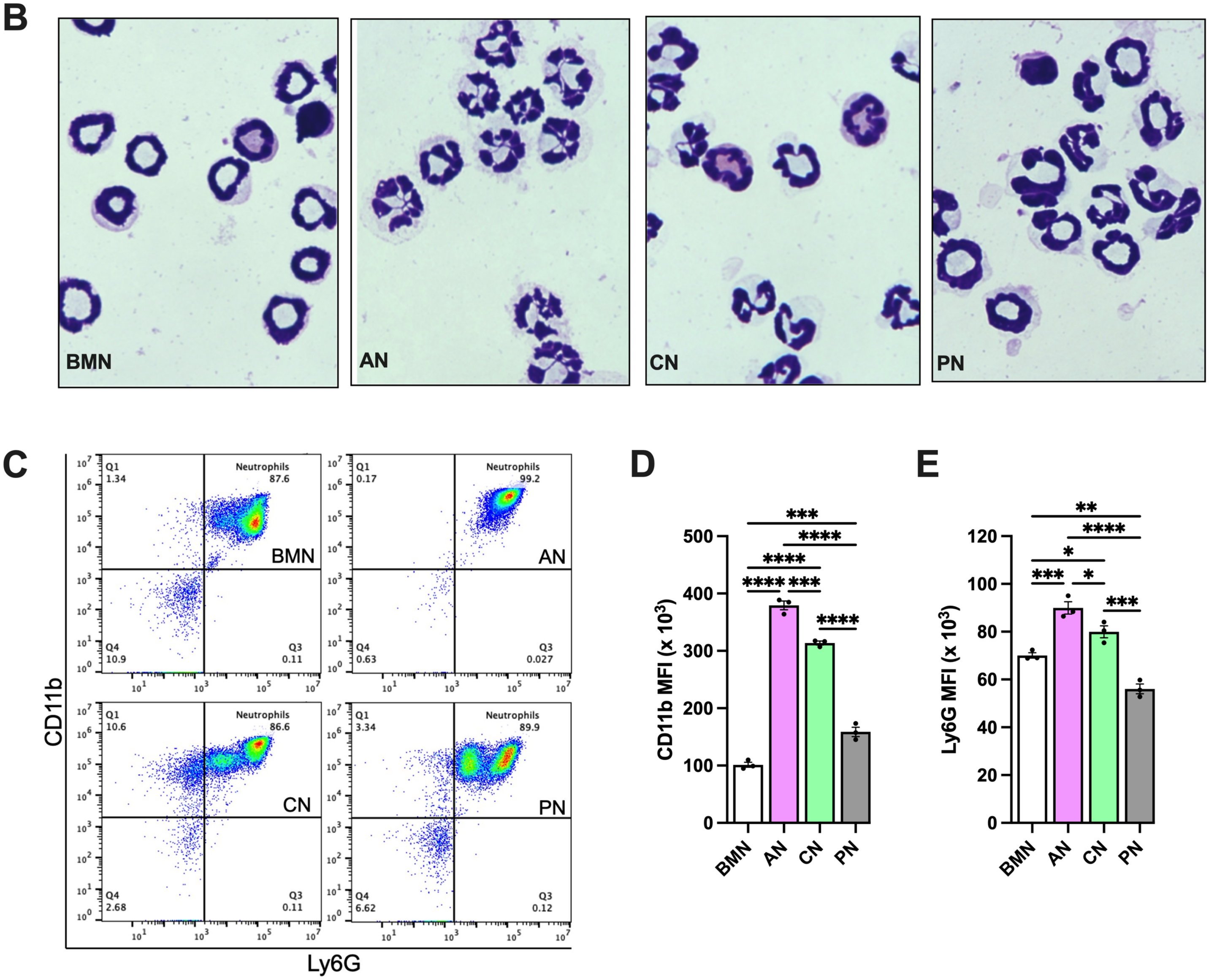


**Supplemental Figure 4 (related to Figure 5)**: **Alveolar neutrophils present more segmented nuclei and mature CD11b and Ly6G expression.** Bone marrow neutrophils (BMN), alveolar neutrophils (AN), circulating neutrophils (CN) and peritoneal neutrophils (PN) were isolated. The total cell number obtained per mouse and the viability are shown in the table (A). Hemacolor-stained cells showing neutrophils nuclei morphology (B). Representative flow cytometry plots of neutrophils, represented as $CD11b^+Ly6G^+$ cells in the $CD45.2^+$ gate (C). Mean fluorescence intensity of CD11b (D) and Ly6G (E) on neutrophils. ****, ***, ** and * indicate $p<0.0001$, $p<0.001$, $p<0.01$ and $p<0.05$, respectively, determined by one-way ANOVA followed by Tukey's multiple comparisons test; values are the mean ± SEM; n=3.

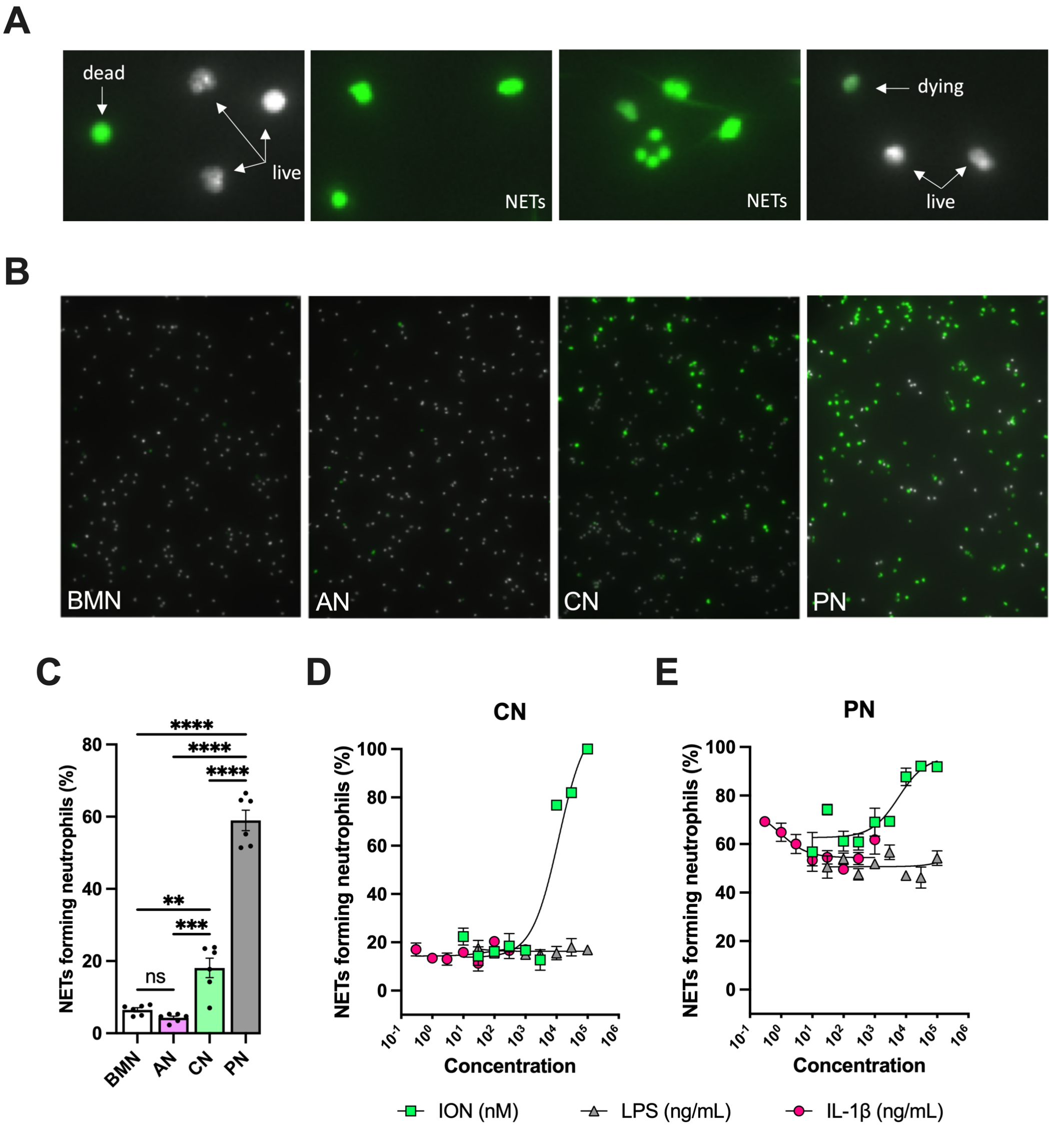


**Supplemental Figure 5 (related to Figure 5)**: **Non-stimulated circulating neutrophils and peritoneal neutrophils are more susceptible to form NETs compared with bone marrow neutrophils and alveolar neutrophils.** NETs are represented by elongated-shape Sytox green-positive (green) Hoechst-negative (white) cells. Sytox green-negative Hoechst-positive cells are considered live neutrophils, and round-shape Sytox green-positive Hoechst-negative cells are classified as dead neutrophils. Dying neutrophils are shown as Sytox green-positive Hoechst-positive cells (A). Hoechst-stained neutrophils were incubated without exogenous stimulations for 5 hours, stained with Sytox green and the images were captured under the microscope. Representative images (A) and NETs forming neutrophils (B) of non-stimulated BMN, AN, CN and PN. CN (C) and PN (D) were stimulated with various concentrations of lipopolysaccharide (LPS, 30-100,000 ng/mL), IL-1β (0.3-1000 ng/mL), and ionomycin (ION) (10-100,000 nM) and the curves of NETs forming neutrophils were plotted. ****, ***, and ** indicate $p<0.0001$, $p<0.001$ and $p<0.01$, respectively, determined by one-way ANOVA followed by Tukey's multiple comparisons test; ns, non-significant; values are the mean ± SEM; representative of 3 independent experiments.

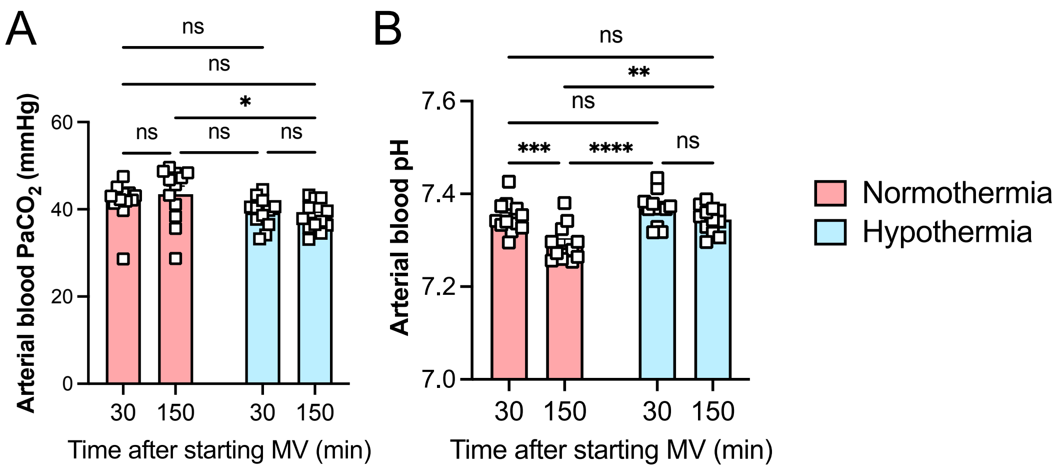


**Supplemental Figure 6 (related to Figure 6): Hypothermia protectes against LPS plus high volume mechanical ventilation-induced severe acute lung injury without respiratory acidosis or alkalosis.** LPS was intratracheally instilled into C57BL/6 mice and, after 120 minutes, the animals were anesthetized and placed on mechanical ventilation (MV) for 180 minutes with HVV. Arterial blood partial pressure of carbon dioxide (arterial blood $PaCO_2$) (A), pH (B) at 30 and 150 min after HVV. *, **, *** and **** were determined by two-way ANOVA followed by Tukey's multiple comparisons test.

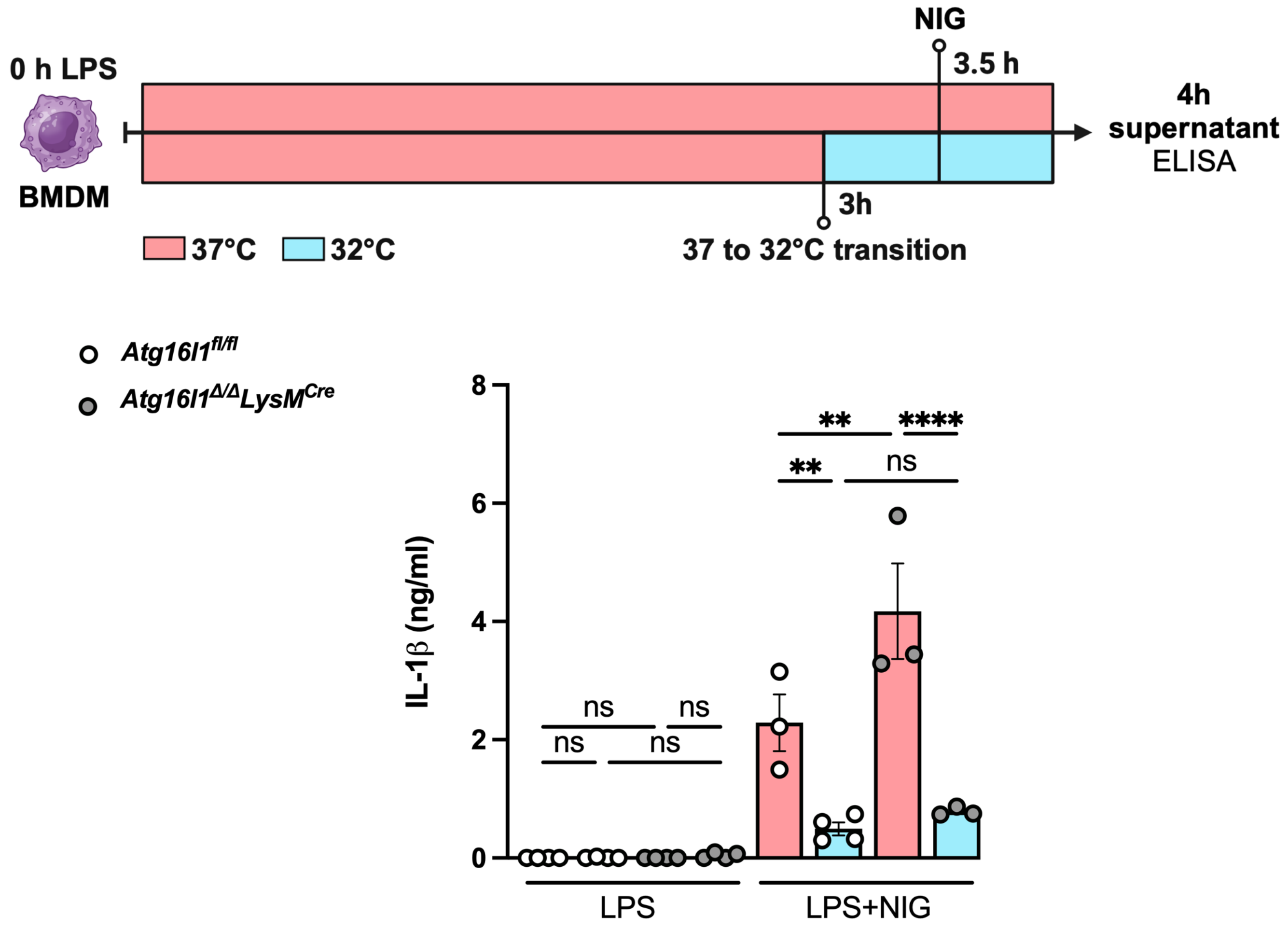


**Supplemental Figure 7 (related to Figure 7): Hypothermia-induced NLRP3 inflammasome inhibition is independent of autophagy.** Bone marrow derived macrophages (BMDM) isolated from *Atg16l1*$^{fl/fl}$ or *Atg16l*$^{\Delta/\Delta}$ *Lysm*$^{Cre}$ were primed with LPS for 3h at 37 ℃, incubated at 37 ℃ or 32 ℃ for 30 min prior NIG treatment for another 30 min. The IL-1β concentration in the culture supernatants was determined by ELISA. ** indicates $p$<0.01 determined by three-way ANOVA followed by Tukey's multiple comparisons test; ns, non-significant; values are the mean ± SEM; representative of 3 independent experiments.

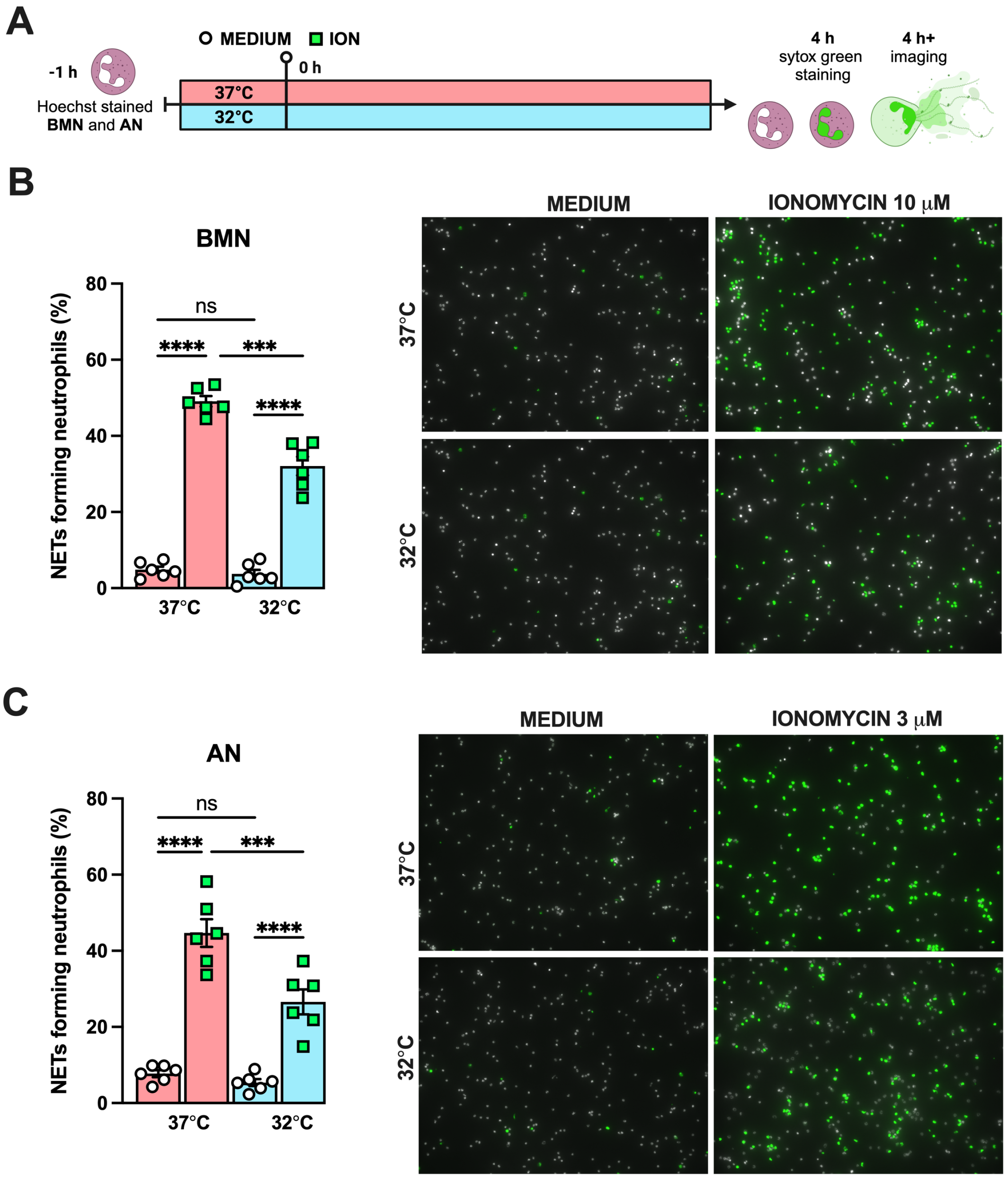


**Supplemental Figure 8 (related to Figure 6): Hypothermia inhibits NETs formation *in vitro.*** Bone marrow neutrophils BMN and alveolar neutrophils AN were incubated for 1 hour prior stimulation at 37°C or 32 °C. Neutrophils were stimulated with ION for 4 hours at their respective temperatures in the presence of Hoechst, then stained with Sytox green and the images were captured (A). BMN were stimulated with 10 μM of ION (B) while AN were stimulated with 3 μM of ION (D). NETs formation was then evaluated. **** and *** indicate $p<0.0001$ and $p<0.001$, respectively, determined by two-way ANOVA followed by Tukey's multiple comparisons test; ns, non-significant; values are the mean ± SEM, representative of 3 independent experiments.
